# Transfer of learned cognitive flexibility to novel stimuli and task sets

**DOI:** 10.1101/2021.07.21.453253

**Authors:** Tanya Wen, Raphael M. Geddert, Seth Madlon-Kay, Tobias Egner

**Affiliations:** Center for Cognitive Neuroscience, Duke University, Durham, NC, USA; Department of Psychology and Neuroscience, Duke University, Durham, NC, USA

**Keywords:** cognitive flexibility, reinforcement learning, generalization, task switching, meta-flexibility

## Abstract

Adaptive behavior requires learning about the structure of one’s environment to derive optimal action policies, and previous studies have documented transfer of such structural knowledge to bias choices in new environments. Here, we asked whether people could also acquire and transfer more abstract knowledge across different task environments, specifically expectations about cognitive control demands. Over three experiments, participants performed a probabilistic card-sorting task in environments of either a low or high volatility of task rule changes (requiring low or high cognitive flexibility respectively) before transitioning to a medium-volatility environment. Using reinforcement learning modeling, we consistently found that previous exposure to high task rule volatilities led to faster adaptation to rule changes in the subsequent transfer phase. These transfers of expectations about cognitive flexibility demands were both task- (Experiment 2) and stimulus- (Experiment 3) independent, thus demonstrating the formation and generalization of environmental structure knowledge to guide cognitive control.

**Statement of Relevance:** We investigated whether structural knowledge of one task environment can be transferred to guide cognitive control strategies in new environments. Past research has found that while learning generally improves subsequent performance, it does so only for the learned task (“near transfer”) and has little or no generalizability to novel task rules and stimuli (“far transfer”). However, recent studies suggest that learning more abstract, structural task features (e.g., cognitive maps) allows for that knowledge to be applied to new environments. Here, we took a critical additional step and showed that people can acquire and transfer expectations about cognitive control demands (specifically cognitive flexibility) across different task environments. To our knowledge, this is the first demonstration of people’s ability to extract and re-use cognitive control learning parameters that transcend specific stimuli and tasks. This transfer of learned cognitive flexibility is particularly noteworthy because such flexibility is impaired in several common psychiatric conditions.

## Introduction

Adaptive behavior requires us to identify and keep in mind the currently relevant “rules of the game” – that is, which responses to which stimuli likely lead to desirable outcomes (also known as task sets; Monsell, 2003). Moreover, given that the world is ever-changing, optimal regulation of task sets involves resolving a tradeoff between needing to implement the current task set and shielding it from distraction (cognitive stability) versus being ready to update (or switch) task sets in response to changing environmental contingencies (cognitive flexibility; Goschke, 2003; Nassar & Troiani, 2020). Importantly, neither stability nor flexibility are inherently beneficial; rather, it is the ability to dynamically adapt one’s flexibility level to suit varying environmental demands, referred to as meta-flexibility, that facilitates optimal cognition (Goschke, 2013).

To adjust cognitive flexibility in an optimal manner, one must infer which task sets to use at a given time by observing environmental statistics (Behrens, Woolrich, Walton, & Rushworth, 2007; Yu, Wilson, & Nassar, 2020), such as associations between stimuli, responses, and outcomes. The process of learning these associations can be characterized by reinforcement learning (RL) models (Barraclough, Conroy, & Lee, 2004; Lee, Seo, & Jung, 2012; Sutton & Barto, 1998). In the context of task switching, the value to be learned is the likelihood that a given task set is currently relevant. The level of cognitive flexibility a learner exhibits can be described by their learning rate, which determines the degree to which recent feedback updates their beliefs. Previous studies have shown that people’s learning rates are typically low during periods of environmental stability and high during periods of volatility (Behrens et al., 2007; Browning, Behrens, Jocham, O’Reilly, & Bishop, 2015; Jiang, Beck, Heller, & Egner, 2015; Jiang, Heller, & Egner, 2014; Massi, Donahue, & Lee, 2018), though those studies have looked at the direct learning of stimulus-reward associations or proportions of stimulus types. To our knowledge, it has not been tested whether environmental volatility can similarly influence the learning of higher-order rules such as task sets.

Moreover, adapting cognitive flexibility is impossible in a previously unobserved environment. How, then, do people set their cognitive flexibility in new situations? We posit that successfully matching cognitive flexibility levels to varying demand contexts could be mediated by learning and transferring knowledge about the demand structure of previous environments to novel environments. For example, while learning a novel task, it may be beneficial to exploit relevant information acquired in the past (Kemp, Goodman, & Tenenbaum, 2010; Mark, Moran, Parr, Kennerley, & Behrens, 2020; Yu et al., 2020). Previous studies have demonstrated that structural knowledge of an environment in the form of cognitive maps of stimulus associations (Mark et al., 2020) and correlated bandit arms (Baram, Muller, Nili, Garvert, & Behrens, 2020; Schulz, Franklin, & Gershman, 2020) can foster transferrable expectations about the structure of new environments. However, to the best of our knowledge, it has not been tested whether learning parameters driving cognitive control processes, such as task set updating, can be transferred to different contexts.

We here combined these two prior insights – volatility learning and structure transfer – to create a novel test of the acquisition and transfer of cognitive control policies, specifically, one’s level of cognitive flexibility or switch-readiness. We hypothesize that, first, cognitive flexibility is adjusted in response to environmental volatility, and second, that a level of cognitive flexibility learned in one environment can be transferred to another. Observing such transfer of learned cognitive flexibility would a novel finding in the fields of decision-making and cognitive control.

To this end, the current study investigated whether participants learning to update task sets faster or slower (i.e., at different learning rates) in one context would transfer their expectations to another context. Specifically, we conducted three experiments employing a probabilistic version of the Wisconsin Card Sorting Task (Berg, 1948; Van Eylen et al., 2011) wherein two groups of participants were initially exposed to either a low- or high-volatility learning environment, with seldom vs. frequent rule changes, respectively. Next, participants from both groups switched to the same medium-volatility transfer environment, which had an intermediate rate of rule changes. RL models were fit to participants’ rule-choice behavior to quantify the rates of rule-updating (i.e., the learning rates) of the two groups in both task phases. We predicted that participants who encountered the low-volatility condition would have lower learning rates than participants who experienced the high-volatility condition, and that these learning rates would generalize to the transfer phase. Across the three experiments, we systematically decreased the task and stimulus overlap to investigate whether the similarity between the learning and transfer phases influenced learning rate transfer.

## General Methods

In three experiments, we examined whether participants could acquire and transfer knowledge about cognitive flexibility demands across different contexts. In all experiments, participants were split into two groups (low- and high-volatility) that completed a learning phase where the task sets switched less or more frequently, respectively. Next, we tested them in a medium-volatility environment transfer phase, where the switch rate of task rules was the same for both groups. Our main question was whether expectations about the frequency of task-set updating acquired in the learning phase would generalize to the subsequent transfer phase.

### Procedure

Figure 1 illustrates two example trials of the task paradigm in the learning phase of all three experiments. On each trial, three cards arranged in a pyramid were simultaneously presented on the screen, randomly chosen with the following constraints. The card on the top served as the reference card, and the cards at the bottom were choice cards. One of the two choice cards shared the same value in one dimension (e.g., shape) as the reference card, but had different values in all the other dimensions (e.g., color, filling, number). The other choice card shared the same value as the reference card in a second dimension (e.g., color), while being different in the three other dimensions (e.g., shape, filling, number). Additionally, there were no shared values on any dimensions between the two choice cards. Only two of the four dimensions, randomly assigned for each participant, were relevant as possible matching rules during the experiment. The two relevant dimensions were explicitly instructed to the participant (and practiced, see below) prior to the experiment. Only one of the two dimensions was the valid matching rule at any one time, and the valid rule changed over time. It was the participants’ goal to figure out, via trial-and-error learning, which matching rule was currently valid on a given trial.

**Figure 1.**
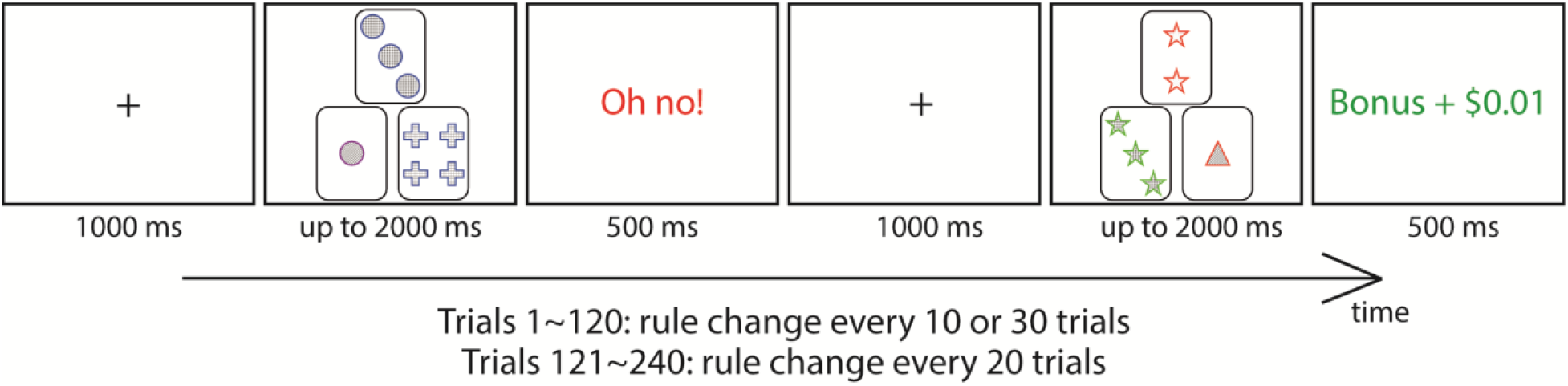
Illustration of the task paradigm. Each trial began with a fixation period, followed by a display of the reference card (top) and two choice cards (bottom) that required a participant response, followed by feedback. Participants were asked to match the correct choice card with the reference card according to the dimension (i.e., color, shape, filling, or number) they believed to be the currently relevant matching rule. In the example above, participants had to sort cards according to color or shape. In the first half of the experiment (learning phase), the sorting rule changed every 30 trials for participants in the low-volatility group, and every 10 trials for participants in the high-volatility group. In the second half of the experiment (transfer phase), the sorting rule changed every 20 trials for both groups.

Each trial began with a 1 s fixation period. Then, participants were asked to match the reference card to the correct choice card based on the dimension they believed to be the currently valid matching rule, using the “z” or “m” button to indicate the left or right choice card, respectively. The cards remained on the screen for up to 2 s or until participants made a response. If participants did not respond in time, they would receive a “Too slow!” feedback and would be asked to press the spacebar to begin the next trial. Otherwise, they would be given either an “Oh no!” or “Bonus + $0.01” feedback for 500 ms. The feedback validity was 80%; that is, participants had an 80% chance of receiving positive feedback (and a 20% chance of negative feedback) on correct responses, and vice versa for incorrect responses. Participants were informed of the 80% feedback validity before the experiment. The correct sorting dimension stayed the same for a fixed number of trials before changing to the other dimension, but participants were not explicitly informed about the frequency of rule changes.

Before starting the main experiment, participants were asked to perform a practice task consisting of 40 trials, with the sorting rule changing after 20 trials. The practice task was similar to the main experiment, except that the sorting rule was explicitly displayed on the screen. Participants had to achieve at least 90% accuracy on the practice task to move on to the main experiment. In the main experiment, both the low- and high-volatility groups completed a total of 240 trials. In the first half of the experiment (the learning phase), the sorting rule changed every 30 trials for participants in the low-volatility group, and every 10 trials for participants in the high-volatility group. In the second half of the experiment (the transfer phase), the sorting rule changed every 20 trials for both groups. In Experiment 1, the stimuli and task sets remained the same during the transfer phase as in the learning phase; in Experiment 2, the stimuli remained the same, but task sets were novel; and in Experiment 3, both stimuli and task sets were novel. There were no explicit instructions informing participants about the currently relevant rule, such that participants always had to rely on the feedback to figure out the currently relevant sorting rule to maximize their earnings.

### Behavioral analyses

For each experiment, we compared the accuracies of the low- and high-volatility groups, with an accurate trial defined as responding according to the correct sorting rule, regardless of feedback. We then split the data into learning and transfer phases, and calculated the accuracies for each phase, and entered participants’ mean accuracy values into a phase (learning vs. transfer) × volatility (low vs. high) ANOVA. This was to ensure that any group differences in RL model parameters were not confounded by differences in overall accuracy.

We hypothesized that participants in the low volatility groups would be slower to switch tasks in response to a rule change than the high volatility participants. To test this directly, we examined the probability of participants choosing the currently (or previously) active rule on trials before (and after) the periodic rule change point. For both learning and transfer phases, we averaged individuals’ choice probability from −5 to +5 trials around the change point. Choice probability was entered into a phase (learning vs. transfer) × boundary (before vs. after) × volatility (low vs. high) ANOVA. Since we hypothesized a priori that the high-volatility group would be quicker to switch after a rule change, we also compared volatility effects separately in each phase and time-bin in planned follow-up analyses.

### Reinforcement-learning modeling

We fit RL models (Sutton & Barto, 1998) to the choice behavior to estimate learning rates for the low- and high-volatility groups in the learning and transfer phases of the three experiments, using Markov Chain Monte Carlo (MCMC) sampling via the “stan” function from the RStan package (Stan Development Team, 2020) in R (R Core Team, 2020). We ran four MCMC chains for 1000 samples, discarding the first 150 as warm-up.

Our first model was a standard hierarchical RL model (RW-RL; Rescorla & Wagner, 1972), fit to the rule choice behavior. A hierarchical model was used as it takes into account the within-subject error of each subject’s parameter estimate, unlike in the classic approach of comparing the mean value of each parameter for each condition after estimating point estimates for each subject (Daw, 2009). The model consisted first of a Q-learning model (Watkins & Dayan, 1992), whereby value estimates for each rule are updated over time based on feedback. Specifically, after an individual (*i*) on trial (*t*) chooses a matching rule, *C_i,t_* ∈ {1, 2} (e.g., color or shape), and feedback is received for that choice, *R_i,t_* ∈ {0, 1} (0 if negative feedback and 1 if positive feedback), the value estimate of that rule, *V*(*C*), is updated according to the following:

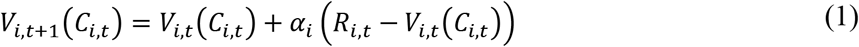

where *α_i_* is each individual’s learning rate. The first trial of each experiment, *V*_*i*1_, as well as the first trial of the transfer phase in Experiments 2 and 3, *V*_*i*,121_, were initialized with a separate starting utility, with the prior distribution of *N*(0.5,0.5). This was not done for the first trial of the transfer phase of Experiment 1 because in that experiment there was no change in stimuli or task between learning and transfer phases.

In order to estimate the distribution of learning rates across experiments, conditions, and individuals, we estimated a multi-level model with three levels of hierarchy. The top level of the hierarchy described how the average learning rate varied across different conditions, while the middle level described how learning rates varied among individuals within a condition. Finally, the bottom level, described by equations (1), (4), and (7), modeled how individuals learned from feedback over the course of the task and predicted their choices. The advantage of using a single hierarchical model across all experiments and conditions is to pool information across conditions, resulting in less noisy estimates and reducing overfitting to individual conditions (Gelman, Hill, & Yajima, 2012).

At the top level of the hierarchy, we assumed the average learning rates for each condition were generated by a mixed-effects general linear model:

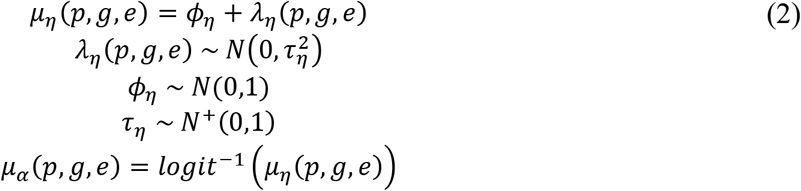

Each condition was defined by a combination of a phase *p* (learning or transfer), a group *g* (high or low volatility), and an experiment *e* (experiment 1, 2, or 3). The hyperparameter *ϕ_η_* is the population average learning rate for all subjects across all conditions. The condition-level random effects *λ_η_*(*p, g, e*) determine how far the average of each condition, i.e., phase (*p*) for each group (*g*) in each experiment (*e*), is from the average of the population mean, with the variance 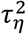 governing the overall variability across conditions.

Next, for the middle level of the hierarchy, we modeled the learning rates of individual participants as arising from a mixed-effects general linear model:

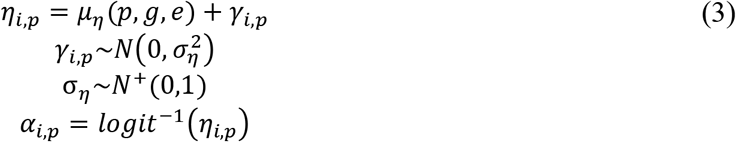

where the random effects *γ_i,p_* determine how far each individual *i* is from the average for their condition, *μ_η_*(*p, g, e*), with 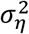 governing the overall variability across subjects within each condition.

We examined the contrasts between conditions of interest. In particular, for each experiment and phase, we compared whether there were any differences in learning rate between the low- and high-volatility groups. We further examined whether this differed across experiments.

We also sought to examine whether the effects of learning rates were differentially driven by positive feedback (rewarded) versus negative feedback (unrewarded) trials. Intuitively, the negative feedback trials would be expected to drive rule switches, as participants would presumably recognize that their currently applied rule was incorrect. We therefore fit a second model, which we call the two-rates (2R) RL model, in which learning rate was fit separately for positive (+) and negative (−) reward (*r*) feedback trials (Donahue and Lee, 2015). The value for each rule, *V*(*C*), is here updated according to:

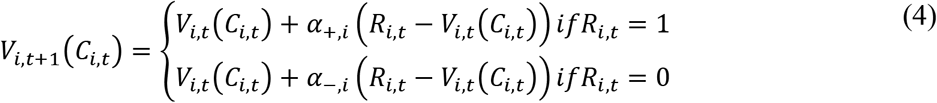

where *α*_+*i*_ is each individual’s learning rate for positive feedback trials, and *α*_−,*i*_ is each individual’s learning rate for negative feedback trials. Similar to the RW-RL model, a hierarchical general linear model is used to estimate mean effects of each condition:

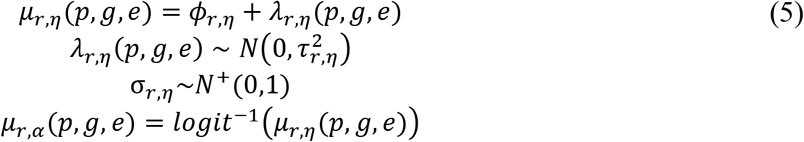

where *ϕ_r,η_* are hyperparameters representing the population mean learning rates for the feedback level *r* for all subjects across the two phases and three experiments. The random effects *λ_r,η_*(*p, g, e*) represent the deviations of each condition from the population means, and 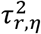 govern the overall variability in each condition.

Another hierarchical general linear model was used to model individual learning rates:

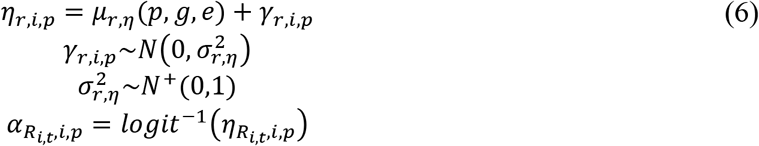

where *μ_r,η_*(*p, g, e*) are hyperparameters representing the mean learning rates for positive and negative feedback trials for each condition, and *γ_r,i,P_* is the deviation of each individual from the condition means, with 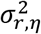 variability.

Finally, we also examined how the two volatility groups differed in terms of their action policies. For both the RW-RL and the 2R-RL model, we assumed that subjects chose rules probabilistically based on the value estimates according to a softmax distribution (Daw 2009). Thus, choice probabilities of selecting each rule (e.g., color or shape) for each trial were computed as follows:

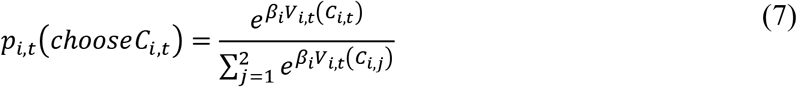

Here, *β_i_* is a hyperparameter known as the inverse temperature, which represents how sensitive choice probabilities are to differences in choice value (Katahira 2015). *β_i_* values were calculated for each subject similar to *η_i_*, with a hyperparameter representing the population’s mean *ϕ_β_*, and how many standard deviations, *λ_β_*(*p, g, e*), each condition deviated from their group’s mean, and then another hyperparameter *μ_β_*(*p, g, e*) representing each condition’s mean, and the deviations of each individual, *γ_i,p_*, from the condition mean.

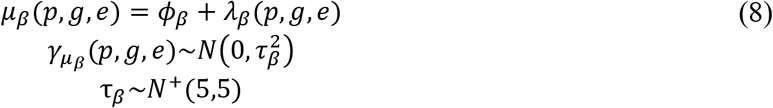

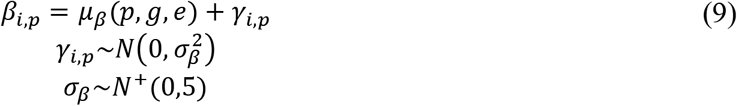

While we report results from both the RW-RL and 2R-RL models, using them to characterize a general overall learning rate (*α_i,p_*) as well as separate learning rates for positive (*α_+,i,p_*) and negative feedback (*α_−,i,p_*) trials, we compared model fits between the two models using the leave-one-out information criterion (Vehtari, Gelman, & Gabry, 2017), and found that the 2R-RL model fit the data better. The expected log pointwise predictive density (ELPD) difference between the two models was −70.1, and its standard error (SE) difference was 30.6. This suggests that in the current experiments, participants did indeed have different learning rates for positive and negative feedback trials.

In the following analyses, we compared the posterior distribution of parameter estimates for learning rates and inverse temperature, in the learning phase and transfer phase, across the three experiments. We report 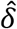, representing the mean difference between conditions in the model. In analyses with multiple factors, the results are reported in the format of an ANOVA. That is, we report the main effect of a factor by comparing the means of each level of that factor, averaging over all other factors. In the case of interactions, we examine the difference of differences between levels in each factor. We also report credible interval (*CI*), which is the Bayesian equivalent of a confidence interval (with a slightly different technical interpretation). All credible intervals reported are central 95% intervals of the posterior differences. Additionally, given our a priori expectation that the learning rate in the high volatility group could only be the same or higher than the low volatility group in the transfer phase, for tests comparing the two volatility groups, we also report 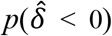, which is the proportion of the posterior difference which falls below zero (corresponding to the logic of a one-tailed *p*-value).

Parameter estimates for the RW-RL model are summarized in Table 1, and for the 2R-RL model in Table 2. Our main analyses focused on learning rates, however. We report the results of inverse temperatures in the Supplementary Results. For visualization, an illustration of learning rate parameter estimates is shown in Figure 4.

**Table 1.**
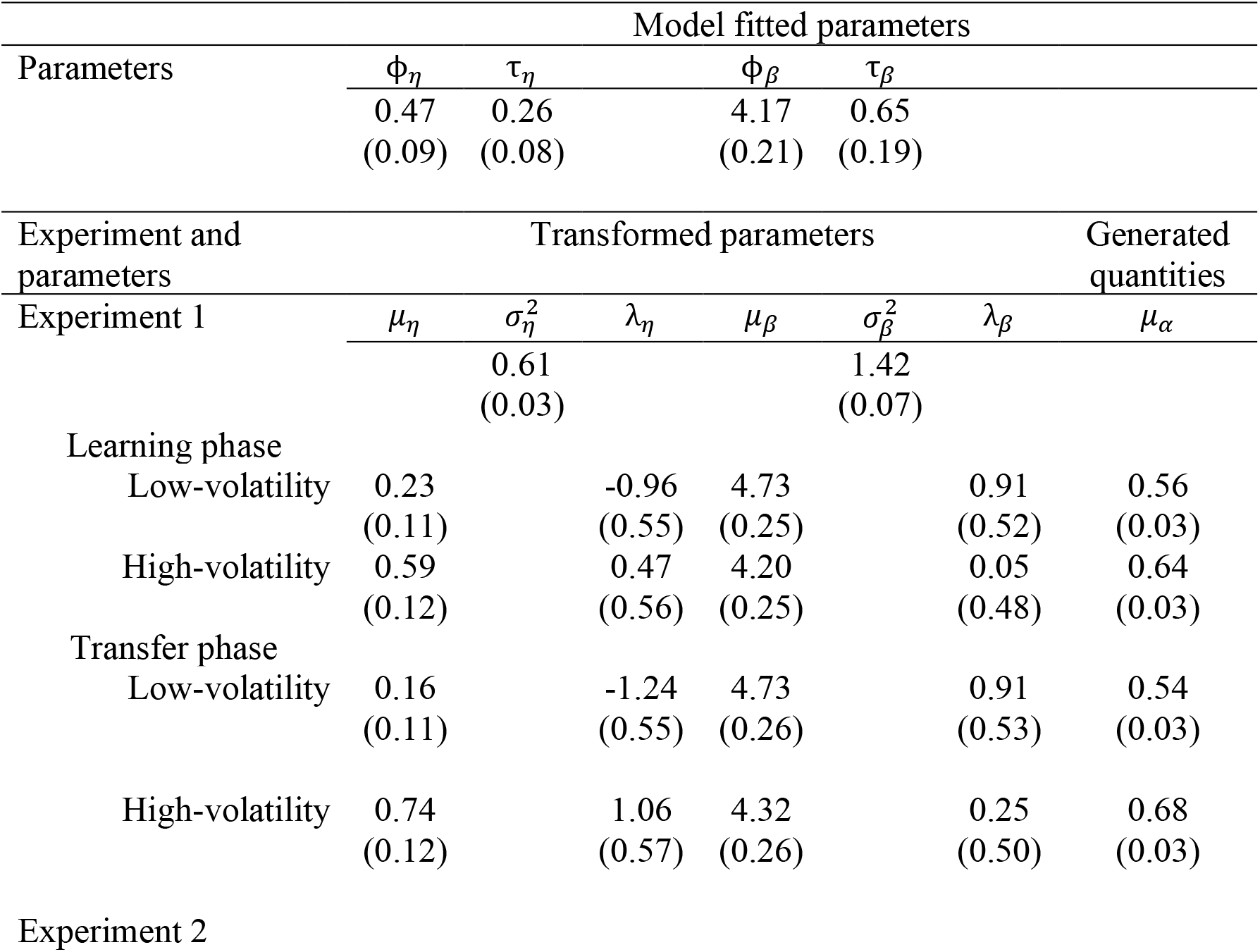
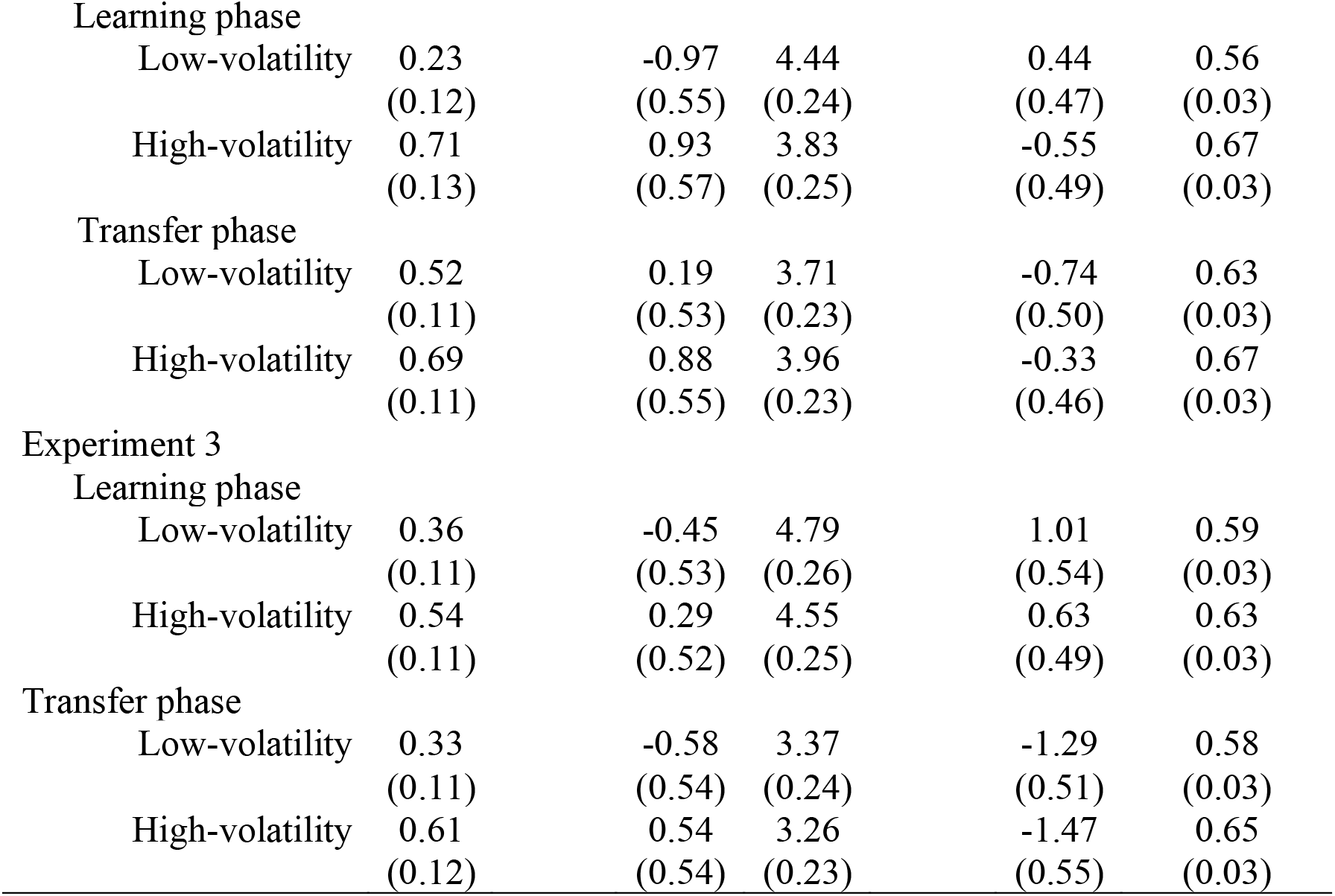
Mean parameter estimates (standard deviations) from the RW-RL model

| Model fitted parameters |  |  |  |  |  |  |  |
| --- | --- | --- | --- | --- | --- | --- | --- |
| Parameters | $\Phi_\eta$ | $\tau_\eta$ | $\Phi_\beta$ | $\tau_\beta$ | | | |
|  | 0.47 | 0.26 | 4.17 | 0.65 |  |  |  |
|  | (0.09) | (0.08) | (0.21) | (0.19) |  |  |  |
| Experiment and parameters | Transformed parameters |  |  |  |  | Generated quantities |  |
| Experiment 1 | $\mu_\eta$ | $\sigma_\eta^2$ | $\lambda_\eta$ | $\mu_\beta$ | $\sigma_\beta^2$ | $\lambda_\beta$ | $\mu_\alpha$ |
|  |  | 0.61 |  |  | 1.42 |  |  |
|  |  | (0.03) |  |  | (0.07) |  |  |
| Learning phase |  |  |  |  |  |  |  |
| Low-volatility | 0.23 |  | -0.96 | 4.73 |  | 0.91 | 0.56 |
|  | (0.11) |  | (0.55) | (0.25) |  | (0.52) | (0.03) |
| High-volatility | 0.59 |  | 0.47 | 4.20 |  | 0.05 | 0.64 |
|  | (0.12) |  | (0.56) | (0.25) |  | (0.48) | (0.03) |
| Transfer phase |  |  |  |  |  |  |  |
| Low-volatility | 0.16 |  | -1.24 | 4.73 |  | 0.91 | 0.54 |
|  | (0.11) |  | (0.55) | (0.26) |  | (0.53) | (0.03) |
| High-volatility | 0.74 |  | 1.06 | 4.32 |  | 0.25 | 0.68 |
|  | (0.12) |  | (0.57) | (0.26) |  | (0.50) | (0.03) |

Experiment 2
|  |  |  |  |  |  |
| --- | --- | --- | --- | --- | --- |
| Learning phase |  |  |  |  |  |
| Low-volatility | 0.23<br>(0.12) | -0.97<br>(0.55) | 4.44<br>(0.24) | 0.44<br>(0.47) | 0.56<br>(0.03) |
| High-volatility | 0.71<br>(0.13) | 0.93<br>(0.57) | 3.83<br>(0.25) | -0.55<br>(0.49) | 0.67<br>(0.03) |
| Transfer phase |  |  |  |  |  |
| Low-volatility | 0.52<br>(0.11) | 0.19<br>(0.53) | 3.71<br>(0.23) | -0.74<br>(0.50) | 0.63<br>(0.03) |
| High-volatility | 0.69<br>(0.11) | 0.88<br>(0.55) | 3.96<br>(0.23) | -0.33<br>(0.46) | 0.67<br>(0.03) |
| Experiment 3 |  |  |  |  |  |
| Learning phase |  |  |  |  |  |
| Low-volatility | 0.36<br>(0.11) | -0.45<br>(0.53) | 4.79<br>(0.26) | 1.01<br>(0.54) | 0.59<br>(0.03) |
| High-volatility | 0.54<br>(0.11) | 0.29<br>(0.52) | 4.55<br>(0.25) | 0.63<br>(0.49) | 0.63<br>(0.03) |
| Transfer phase |  |  |  |  |  |
| Low-volatility | 0.33<br>(0.11) | -0.58<br>(0.54) | 3.37<br>(0.24) | -1.29<br>(0.51) | 0.58<br>(0.03) |
| High-volatility | 0.61<br>(0.12) | 0.54<br>(0.54) | 3.26<br>(0.23) | -1.47<br>(0.55) | 0.65<br>(0.03) |

**Table 2.**
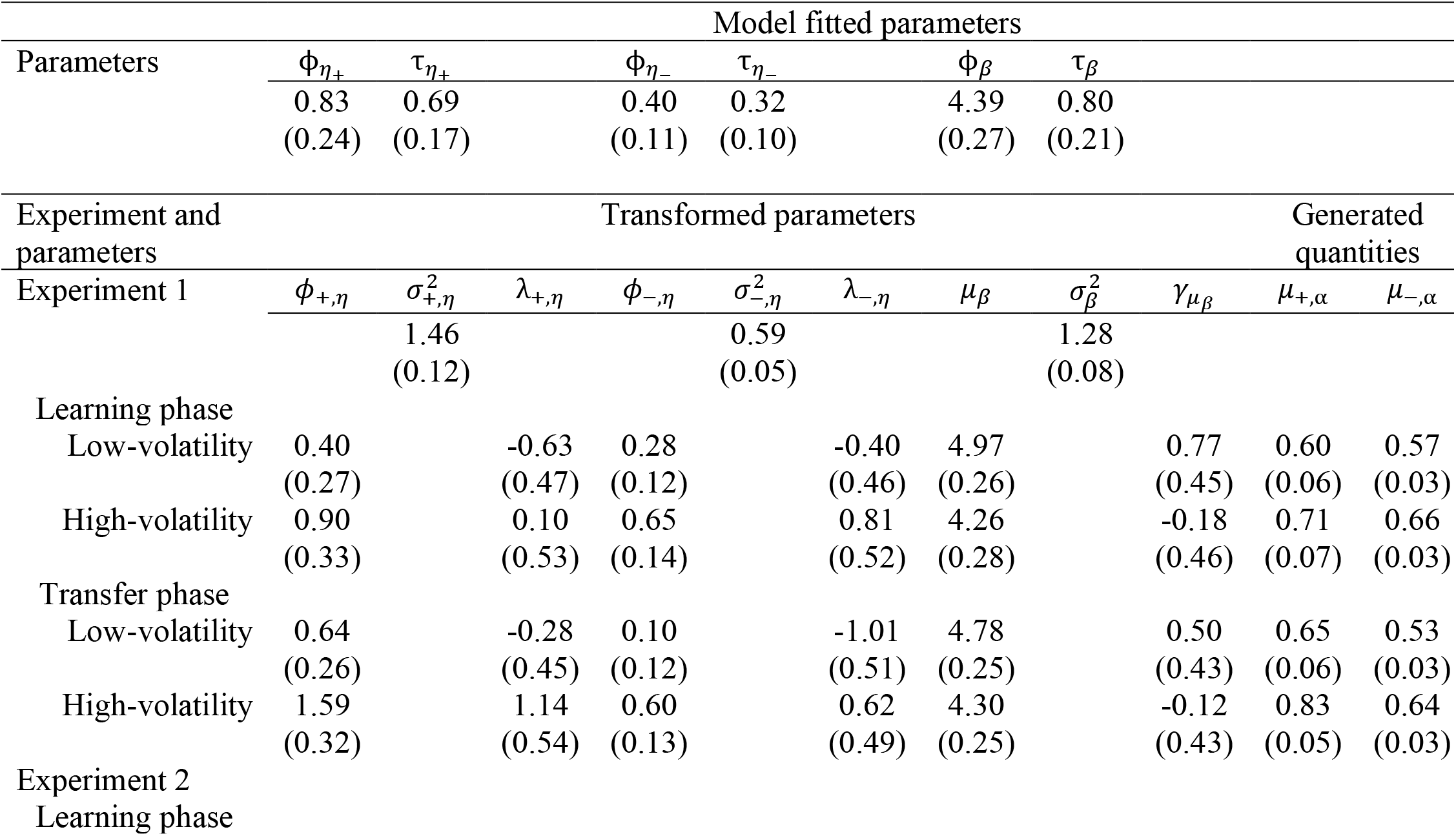
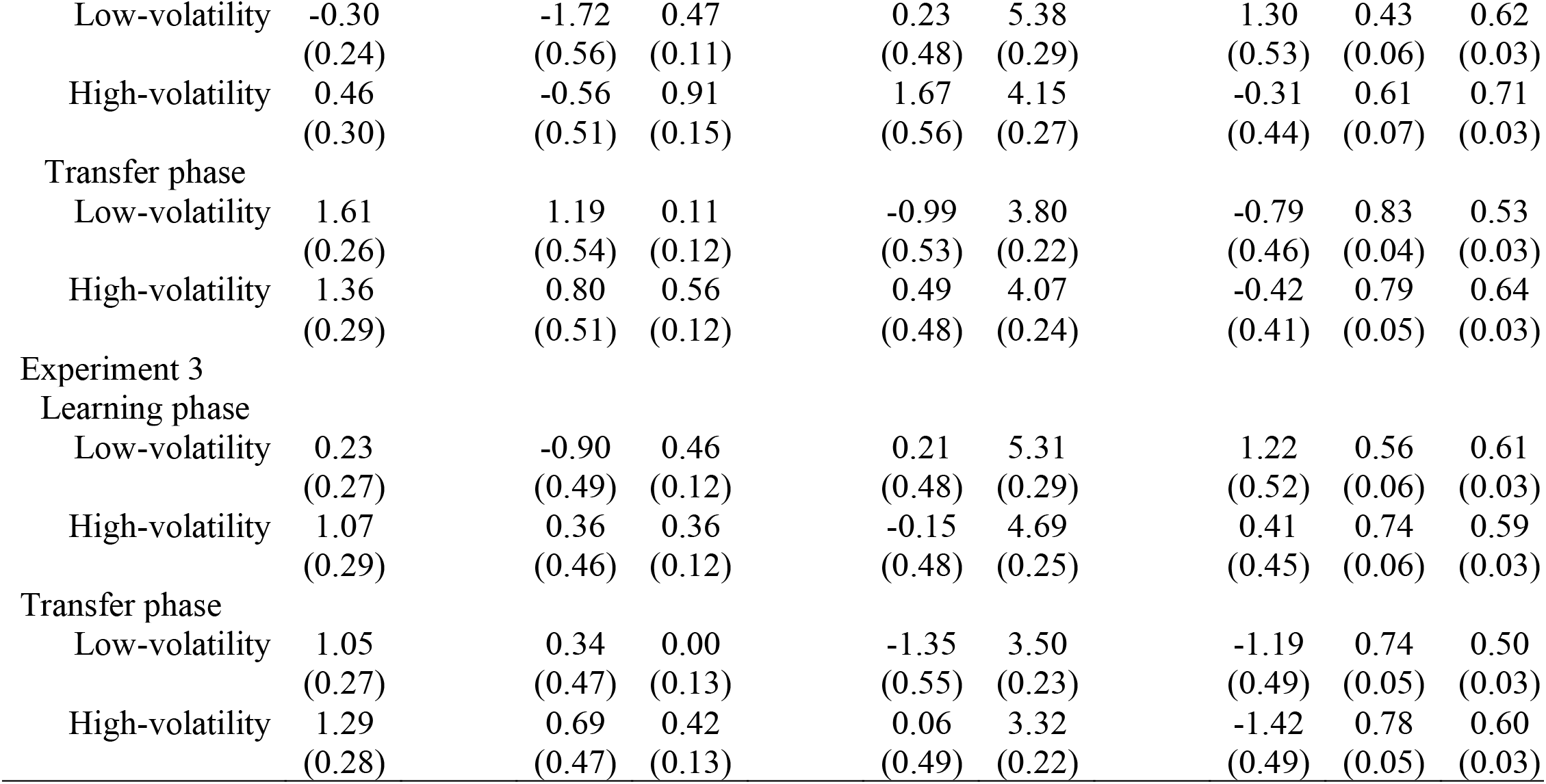
Mean parameter estimates (standard deviations) from the 2R-RL model

| Model fitted parameters |  |  |  |  |  |  |  |  |  |  |  |
| --- | --- | --- | --- | --- | --- | --- | --- | --- | --- | --- | --- |
| Parameters | $\phi_{\eta+}$ | $\tau_{\eta+}$ | | $\phi_{\eta-}$ | $\tau_{\eta-}$ | | $\phi_{\beta}$ | $\tau_{\beta}$ | | | |
|  | 0.83 | 0.69 |  | 0.40 | 0.32 |  | 4.39 | 0.80 |  |  |  |
|  | (0.24) | (0.17) |  | (0.11) | (0.10) |  | (0.27) | (0.21) |  |  |  |
| Experiment and parameters | Transformed parameters |  |  |  |  |  |  |  |  | Generated quantities |  |
| Experiment 1 | $\phi_{+, \eta}$ | $\sigma_{+, \eta}^2$ | $\lambda_{+, \eta}$ | $\phi_{-, \eta}$ | $\sigma_{-, \eta}^2$ | $\lambda_{-, \eta}$ | $\mu_{\beta}$ | $\sigma_{\beta}^2$ | $\gamma_{\mu_{\beta}}$ | $\mu_{+, \alpha}$ | $\mu_{-, \alpha}$ |
|  |  | 1.46 |  |  | 0.59 |  |  | 1.28 |  |  |  |
|  |  | (0.12) |  |  | (0.05) |  |  | (0.08) |  |  |  |
| Learning phase |  |  |  |  |  |  |  |  |  |  |  |
| Low-volatility | 0.40 |  | -0.63 | 0.28 |  | -0.40 | 4.97 |  | 0.77 | 0.60 | 0.57 |
|  | (0.27) |  | (0.47) | (0.12) |  | (0.46) | (0.26) |  | (0.45) | (0.06) | (0.03) |
| High-volatility | 0.90 |  | 0.10 | 0.65 |  | 0.81 | 4.26 |  | -0.18 | 0.71 | 0.66 |
|  | (0.33) |  | (0.53) | (0.14) |  | (0.52) | (0.28) |  | (0.46) | (0.07) | (0.03) |
| Transfer phase |  |  |  |  |  |  |  |  |  |  |  |
| Low-volatility | 0.64 |  | -0.28 | 0.10 |  | -1.01 | 4.78 |  | 0.50 | 0.65 | 0.53 |
|  | (0.26) |  | (0.45) | (0.12) |  | (0.51) | (0.25) |  | (0.43) | (0.06) | (0.03) |
| High-volatility | 1.59 |  | 1.14 | 0.60 |  | 0.62 | 4.30 |  | -0.12 | 0.83 | 0.64 |
|  | (0.32) |  | (0.54) | (0.13) |  | (0.49) | (0.25) |  | (0.43) | (0.05) | (0.03) |
| Experiment 2 |  |  |  |  |  |  |  |  |  |  |  |
| Learning phase |  |  |  |  |  |  |  |  |  |  |  |

|  |  |  |  |  |  |  |  |  |
| --- | --- | --- | --- | --- | --- | --- | --- | --- |
| Low-volatility | -0.30<br>(0.24) | -1.72<br>(0.56) | 0.47<br>(0.11) | 0.23<br>(0.48) | 5.38<br>(0.29) | 1.30<br>(0.53) | 0.43<br>(0.06) | 0.62<br>(0.03) |
| High-volatility | 0.46<br>(0.30) | -0.56<br>(0.51) | 0.91<br>(0.15) | 1.67<br>(0.56) | 4.15<br>(0.27) | -0.31<br>(0.44) | 0.61<br>(0.07) | 0.71<br>(0.03) |
| Transfer phase |  |  |  |  |  |  |  |  |
| Low-volatility | 1.61<br>(0.26) | 1.19<br>(0.54) | 0.11<br>(0.12) | -0.99<br>(0.53) | 3.80<br>(0.22) | -0.79<br>(0.46) | 0.83<br>(0.04) | 0.53<br>(0.03) |
| High-volatility | 1.36<br>(0.29) | 0.80<br>(0.51) | 0.56<br>(0.12) | 0.49<br>(0.48) | 4.07<br>(0.24) | -0.42<br>(0.41) | 0.79<br>(0.05) | 0.64<br>(0.03) |
| Experiment 3 |  |  |  |  |  |  |  |  |
| Learning phase |  |  |  |  |  |  |  |  |
| Low-volatility | 0.23<br>(0.27) | -0.90<br>(0.49) | 0.46<br>(0.12) | 0.21<br>(0.48) | 5.31<br>(0.29) | 1.22<br>(0.52) | 0.56<br>(0.06) | 0.61<br>(0.03) |
| High-volatility | 1.07<br>(0.29) | 0.36<br>(0.46) | 0.36<br>(0.12) | -0.15<br>(0.48) | 4.69<br>(0.25) | 0.41<br>(0.45) | 0.74<br>(0.06) | 0.59<br>(0.03) |
| Transfer phase |  |  |  |  |  |  |  |  |
| Low-volatility | 1.05<br>(0.27) | 0.34<br>(0.47) | 0.00<br>(0.13) | -1.35<br>(0.55) | 3.50<br>(0.23) | -1.19<br>(0.49) | 0.74<br>(0.05) | 0.50<br>(0.03) |
| High-volatility | 1.29<br>(0.28) | 0.69<br>(0.47) | 0.42<br>(0.13) | 0.06<br>(0.49) | 3.32<br>(0.22) | -1.42<br>(0.49) | 0.78<br>(0.05) | 0.60<br>(0.03) |

Finally, we performed a parameter recovery analysis that demonstrates that our models can faithfully reproduce parameter estimates from simulated data. We demonstrate that our models identify differences in learning rates between groups when differences exist, and not when differences do not exist, as our results critically depend on trusting in the group learning rate differences determined by our models. The details of the parameter recovery analysis are reported in the Supplementary Results.

### Data and code sharing

All data and code of experiments and analyses have been deposited on GitHub, https://github.com/tanya-wen/Meta-flexibility.

## Experiment 1

### Method

Experiment 1 examined whether prior exposure to low-vs. high-volatility rule switching environments biased people’s propensity to infer rule changes in response to negative feedback in subsequent medium-volatility environments. More specifically, it explored the transfer of learning rates between initial and subsequent environments that differed solely in terms of rule change volatility, with task stimuli and categorization rules held constant. Later experiments then addressed the question of whether transfer occurs when task rules and stimuli also changed.

#### Participants

Due to a lack of comparable prior studies, we could not base our target sample size on an empirical effect size. We therefore opted for a relatively large target sample size (N=~80). Eighty-eight participants were recruited from Amazon Mechanical Turk (MTurk) and randomly assigned to one of two experimental groups. Participants were compensated at a base pay rate of $2.50 plus any additional bonuses (mean = $1.86, SD = $0.10) earned during the experiment. Thirteen participants were excluded from the analysis due to an overall accuracy lower than 65%, leaving a final sample size of 75. The low-volatility group had 39 participants (22 male, 15 female, 2 did not wish to reply; age range: 26-56, mean = 36.69, SD = 8.81) and the high-volatility group had 36 participants (24 male, 11 female, 1 did not wish to reply; age range: 22-60, mean = 39.44, SD = 10.40).

#### Stimuli

Task stimuli consisted of 256 unique “cards” with one to four display items consisting of a specific shape (circle, triangle, plus, or star) in a particular color (blue, green, red, or purple) and a particular filling (checkered, dots, wave, or grid). We refer to these card properties as “dimensions” (number, shape, color, filling) that can take particular “values” (e.g., 1, 2, 3, or 4 for the number dimension). Each trial involved a display of three such cards. See Figure 1 for example stimuli.

#### Procedure

In the first half of the experiment, participants had to sort cards according to two of the four dimensions (e.g., color and shape; randomly assigned across participants). Sorting rules alternated every 30 trials for the low-volatility group and every 10 trials for the high-volatility group. In the second half (transfer phase) of the experiment, sorting rules alternated every 20 trials in the transfer phase. There was no explicit separation between the first and second half of the experiment.

### Results

#### Behavior

Figure 2 illustrates the rule sequence and choice data from a representative participant from each group. In spite of the 80%-validity probabilistic feedback, participants were able to track the correct rule most of the time (low-volatility group: mean = 79.28%, SD = 5.76%; high-volatility group: mean = 73.19%, SD = 4.13%). A phase (learning vs. transfer) × volatility (low vs. high) ANOVA showed a significant main effect of phase (*F*(1,73) = 39.82, *p* < 0.001), group (*F*(1,73) = 27.27,*p* < 0.001), as well as phase × volatility interaction (*F*(1,73) = 99.74, *p* < 0.001). The main effect of phase was driven by participants having a higher accuracy for the transfer compared to the learning phase (*t* = 3.56, *p* < 0.001), presumably due to a practice effect. The main effect of volatility was driven by the low-volatility group having an overall higher accuracy than the high-volatility group (*t* = 5.22,*p* < 0.001), and this difference was significant in the learning phase (*t* = 11.03, *p* < 0.001), but not in the transfer phase (*t* = −0.99, *p* = 0.33). This effect was expected, given the greater number of (error-inducing) rule reversals in the high-volatility group’s learning phase.

**Figure 2.**
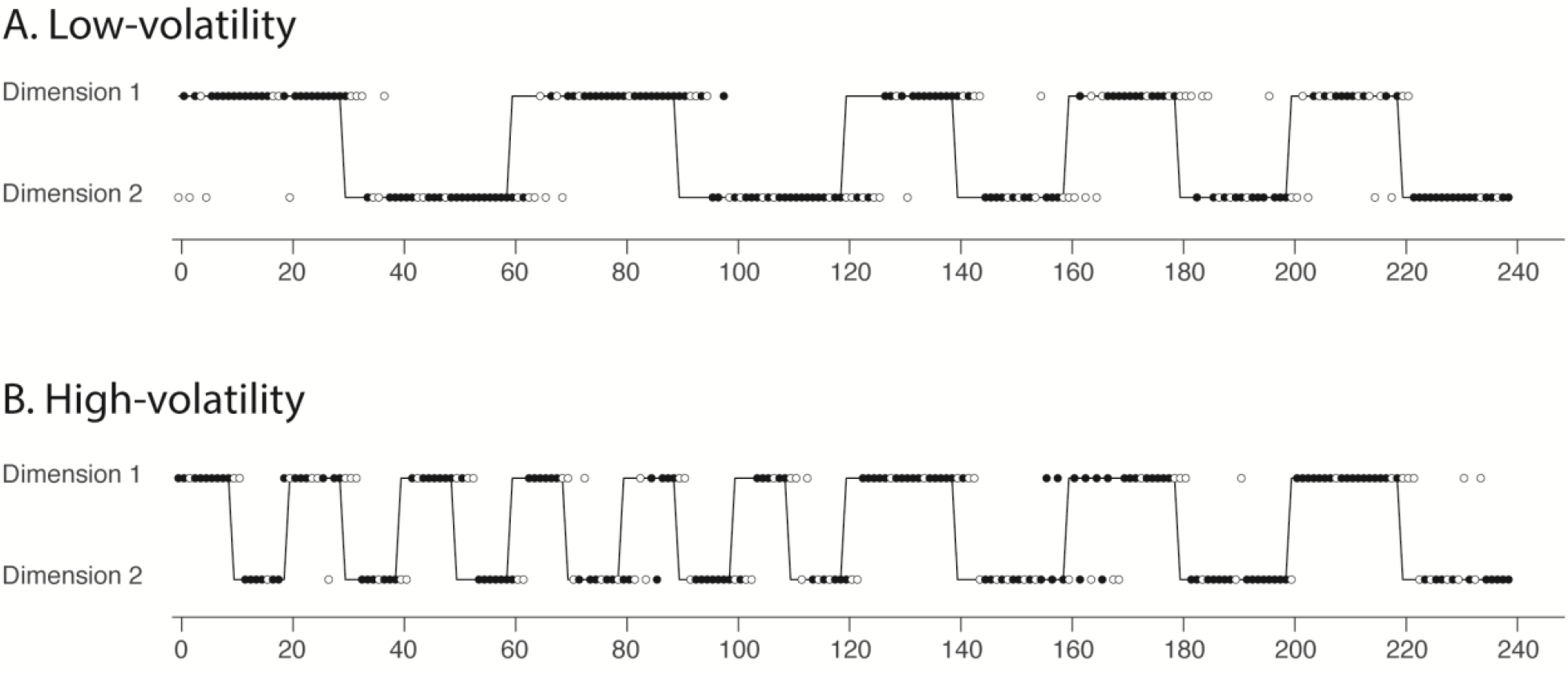
Dimension rule sequences and a representative participant from (A) the low-volatility group and (B) the high-volatility group. On each trial, participants chose a card based on their belief of the currently valid dimensional matching rule, here called Dimension 1 or 2 (circles). They received positive feedback (filled circles) when sorting according to the correct dimension (black line) 80% of the time and for incorrect choices 20% of the time; and they received negative feedback (open circles) for correct choices 20% of the time and for incorrect choices 80% of the time.

Figure 3 illustrates participants’ choice probability as a function of time. We examined whether participants were more likely to switch task rules after the change point. A phase (learning vs. transfer) × boundary (before vs. after) × volatility (low vs. high) ANOVA showed a significant main effect of phase (F(1,73) = 6.76, p = 0.01), a main effect of boundary (F(1,73) = 600.71, p < 0.001), and a main effect of volatility (F(1,73) = 12.57, p < 0.001). There was a phase × boundary interaction (F(1,73) = 20.77, p < 0.001) and a boundary × volatility interaction (F(1,73) = 5.09, p = 0.03). No other interaction was significant. We next conducted separate boundary (before vs. after) × volatility (low vs. high) ANOVAs on the learning and transfer phases. In the learning phase, we found a main effect of boundary (F(1,73) = 225.21, p < 0.001), driven by a higher probability to perform the current task rule prior the rule change. There was also a main effect of volatility (F(1,73) = 10.73, p < 0.01), driven by the low-volatility group being overall more likely to perform the initial task rule. There was no boundary × volatility interaction (F(1,73) = 2.98, p = 0.09). T-tests between volatility groups showed that prior to the rule change, the low-and high-volatility groups were equally likely to perform the current task rule (t = 1.81, p = 0.07). However, the low-volatility group was more likely to persist with the previous rule after the change point (t = 2.93, p < 0.01). In the transfer phase, we found a main effect of boundary (F(1,73) = 693.90, p < 0.001), driven by a higher probability to perform the current task rule prior the rule change. There was also a main effect of volatility (F(1,73) = 6.21, p = 0.02), which was driven by the low-volatility group being overall more likely to perform the initial task rule. There was a marginally significant boundary × volatility interaction (F(1,73) = 3.86, p = 0.05). T-tests between volatility groups showed that prior to the rule change, the low-and high-volatility groups were equally likely to perform the current task rule (t = 0.92, p = 0.36). However, the low-volatility group was more likely to persist with the previous rule after the change point (t = 2.67, p < 0.01).

**Figure 3.**
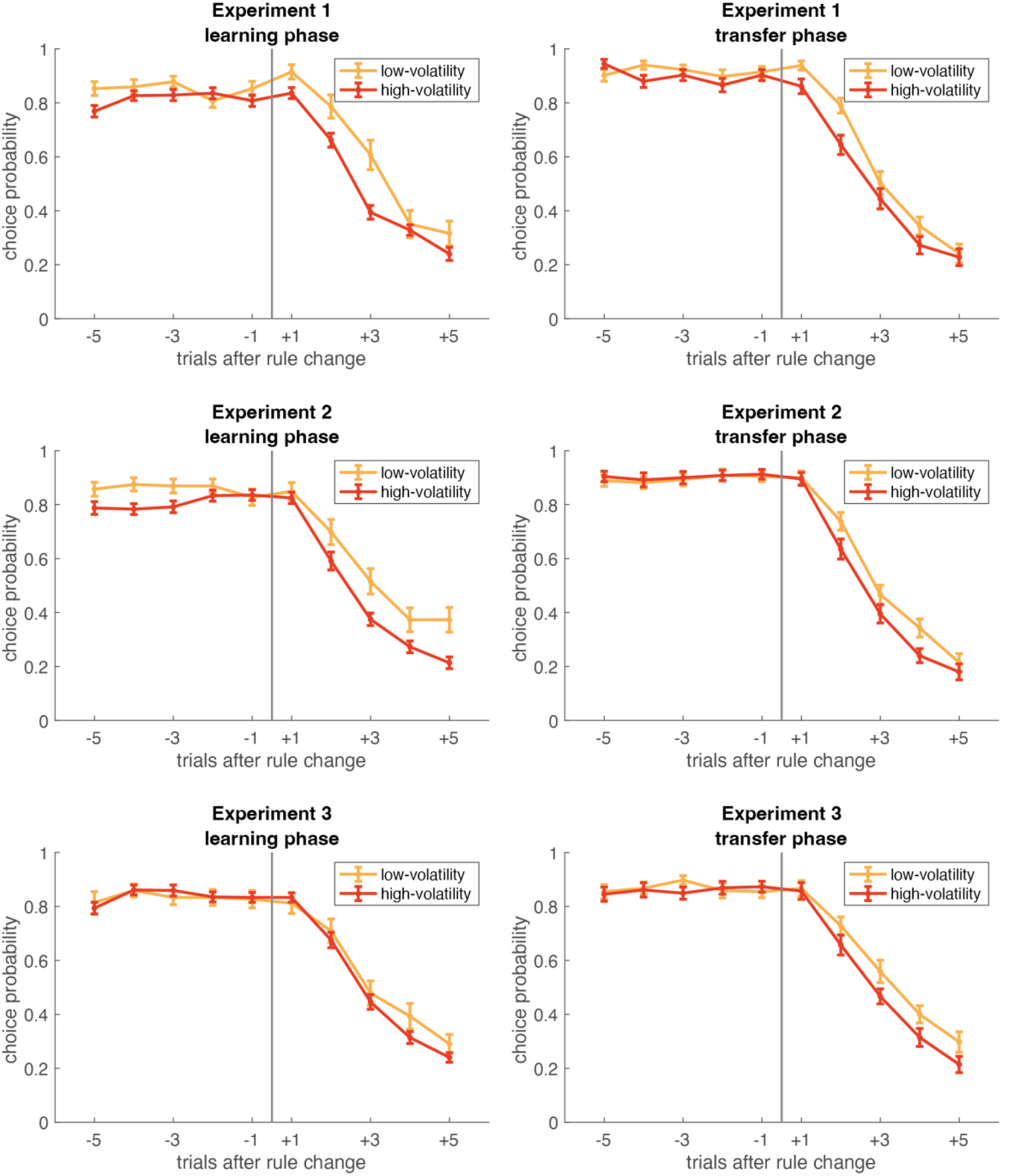
Choice probability as a function of trial before and after the rule change point for the learning and transfer phases of each experiment. Error bars represent standard error.

**Figure 4.**
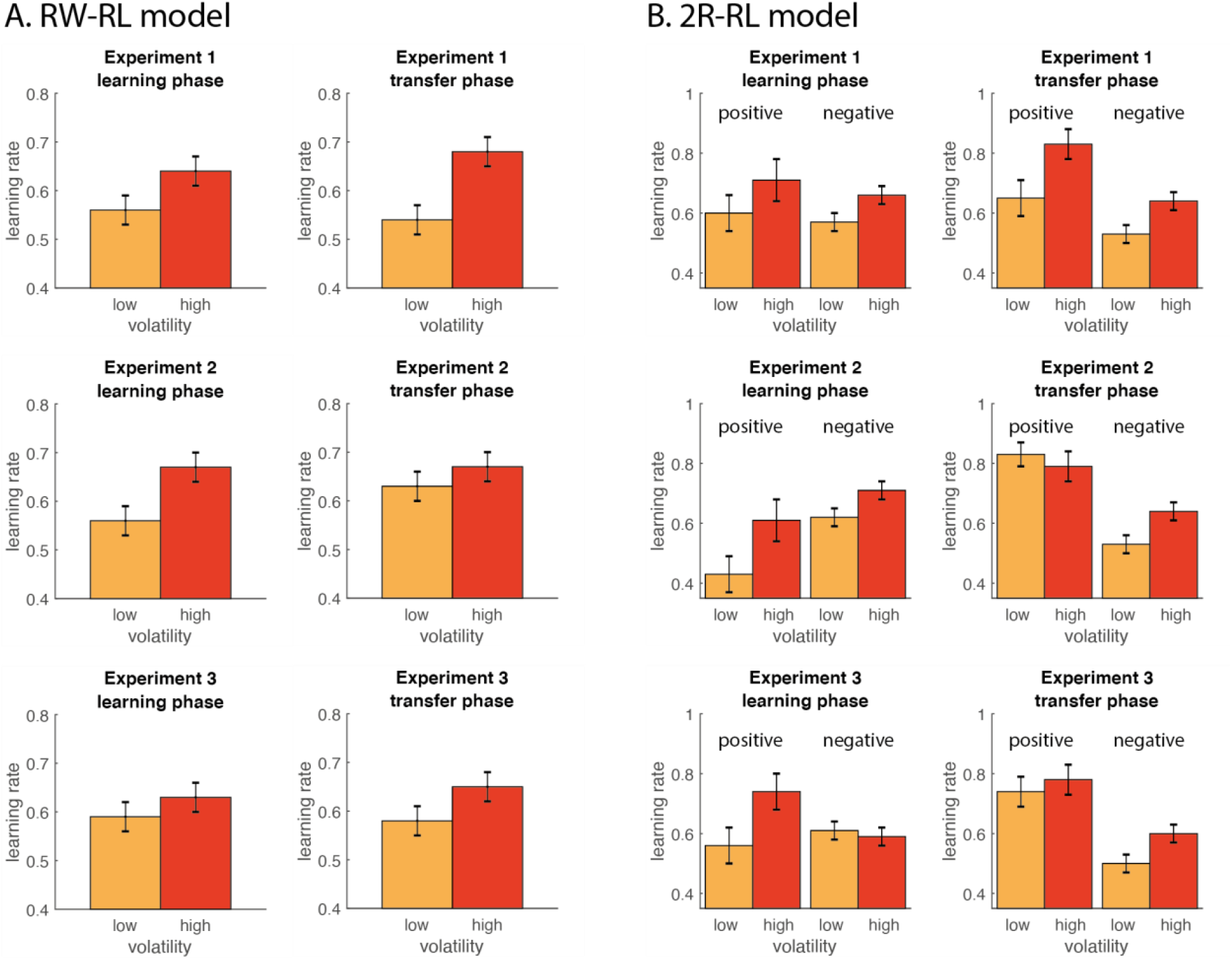
Group level (high versus low volatility) learning rate parameter estimates from (A) the RW-RL model, and (B) the 2R-RL model for the learning and transfer phases in all three experiments. The error bars reflect the fitted group level standard deviation estimates.

#### Reinforcement-learning models

As a confirmatory analysis, we first tested whether learning rates in the high-volatility group were higher than in the low-volatility group during the learning phase, where the matching rule switched every 10 compared to 30 trials. As expected, the RW-RL model showed that learning rates for participants in the high-volatility group were significantly larger than the low-volatility group 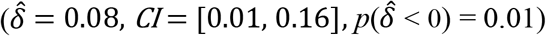. Similarly, in the 2R-RL model, we also found a main effect of volatility, with the high-volatility group exhibiting higher learning rates than the low-volatility group 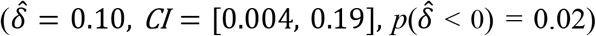. There was no main effect of feedback 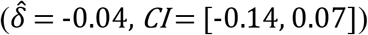, and no feedback × volatility interaction 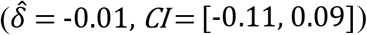.

Our main interest centered on the learning rates during the transfer phase. We found that according to the RW-RL model, the high-volatility group continued to show a higher learning rate than the low-volatility group during this phase 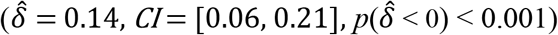. The 2R-RL model showed main effects of volatility 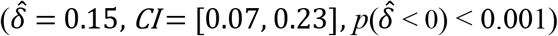, driven by higher learning rates for the high-volatility group; there was also a main effect of feedback 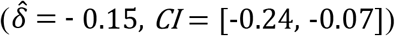, driven by the learning rates being higher for positive feedback trials; and no feedback × volatility interaction 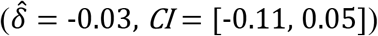. These results document that volatility-driven learning rates acquired during the first half of the task generalized to the second half transfer phase where volatility was equated between groups.

Our results showed that participants adapted their rule switching strategies to the volatility of the task environment, as reflected in faster rule switches around task boundaries, as well as in a higher learning rate in participants in the high-compared to the low-volatility environment. Importantly pre-exposure to high-compared to low-volatility environments led to a higher learning rate in a subsequent medium-volatility environment. In other words, learned expectations about the level of cognitive flexibility required in the environment endured over time.

## Experiment 2

### Method

The results of Experiment 1 suggest that people transfer expectations about the volatility of task rules across adjacent episodes in time (even when the underlying volatility changes) under conditions of identical stimuli and task rules. This represents a temporal “near transfer” of rule learning rate. It is possible that the transfer effect observed in Experiment 1 was driven by participants forming associations between periodic rule switches and the specific rule representations they were learning (see Siqi-Liu & Egner, 2020) rather than reflecting learning at a more abstract level, of making inferences about the general rate at which rules seem to change. Experiment 2 therefore tested whether learning rates would still transfer when the sets of rules that participants learned about changed between the learning and transfer phases (while the stimuli remained the same). Specifically, we probed whether exposure to low- or high-volatility learning environments involving two of four possible task rules (e.g., shape and color matching) would bias the learning rate in subsequent medium-volatility environments with the other two possible task rules (i.e., number and filling matching). Obtaining transfer under these conditions would indicate that the expectations about the rate of rule changes that are being transferred are independent of the specific task rules, thus representing a form of meta-learning.

#### Participants

Ninety-four participants were recruited from MTurk and randomly assigned to one of the two volatility groups. Participants were compensated at a base pay rate of $2.50 plus any additional bonuses (mean = $1.86, SD = $0.10) earned during the experiment. Twelve participants were excluded from the analysis due to overall accuracy lower than 65%, leaving a final sample size of 82. The low-volatility group had 42 participants (20 male, 22 female; age range: 23-70, mean = 39.64, SD = 11.80) and the high-volatility group had 40 participants (22 male, 18 female; age range: 24-76, mean = 38.80, SD = 11.36).

#### Stimuli

The stimuli were the same as in Experiment 1.

#### Procedure

As in Experiment 1, in the first half of the experiment, participants had to sort cards according to two of the four dimensions (e.g., color and shape; randomly assigned across participants). However, unlike Experiment 1, before the start of the second half (transfer phase) of the experiment, participants were taken to another instruction screen and informed that they would now be sorting cards according to the other two dimensions that were previously irrelevant in the first half (e.g., filling and number). There was no practice for the transfer phase, and it started as soon as participants indicated they were ready.

### Results

#### Behavior

Participants were able to perform the task reasonably well (low-volatility group: mean = 79.22%, SD = 5.66%; high-volatility group: mean = 73.58%, SD = 4.45%). The phase (learning vs. transfer) × volatility (low vs. high) ANOVA showed a significant main effect of phase (*F*(1,80) = 22.80, *p* < 0.001), a main effect of volatility (*F*(1,80) = 24.89, *p* < 0.001), and a phase × volatility interaction (*F*(1,80) = 40.93, *p* < 0.001). As in Experiment 1, the main effect of phase was driven by participants having a higher accuracy for the transfer compared to the learning phase (*t* = 4.99, *p* < 0.001), presumably due to generic task practice effects. The main effect of volatility was driven by the low-volatility group having a higher accuracy than the high-volatility group (*t* = 3.71,*p* < 0.001). The interaction was again driven by the low-volatility group having higher accuracy compared to the high-volatility group in the learning (*t* = 8.39, *p* < 0.001), but not in the transfer phase (*t* = −0.54, *p* = 0.59).

We next examined participants’ choice probabilities before and after the rule change point. The phase (learning vs. transfer) × boundary (before vs. after) × volatility (low vs. high) ANOVA showed a significant effect of phase (F(1,80) = 5.67, p = 0.02), a main effect of boundary (F(1,80) = 837.97, p < 0.001), and a main effect of volatility (F(1,80) = 11.52, p = 0.001). There was a phase × boundary interaction (F(1,80) = 10.59, p < 0.01), a phase × volatility interaction (F(1,80) = 4.55, p = 0.04), and a boundary × volatility interaction (F(1,80) = 6.23, p = 0.01). The phase × boundary × volatility interaction was not significant (F(1,80) = 0.16, p = 0.69). We next conducted separate boundary (before vs. after) × volatility (low vs. high) ANOVAs on the learning and transfer phases. In the learning phase, we found a main effect of boundary (F(1,80) = 287.38, p < 0.001)), driven by a higher probability to perform the current task rule prior the rule change. There was a main effect of volatility (F(1,80) = 12.45, p < 0.001), driven by the low volatility group being overall more likely to perform the initial task rule. There was no boundary × volatility interaction (F(1,80) = 1.93, p = 0.17). T-tests between volatility groups showed that prior to the rule change, the low-volatility group was more likely to perform the current task rule (t = 2.37, p = 0.02). Additionally, the low-volatility group was more likely to persist with the previous rule after the change point (t = 3.01, p < 0.01). In the transfer phase, results show a main effect of boundary (F(1,80) = 766.71, p < 0.001), driven by a higher probability to perform the current task rule prior the rule change. There was no main effect of volatility (F(1,80) = 2.65, p = 0.11). There was a significant boundary × volatility interaction (F(1,80) = 6.16, p = 0.02). T-tests between volatility groups showed that prior to the rule change, the low-and high-volatility groups were equally likely to perform the current task rule (t = −0.38, p = 0.71). However, the low-volatility group was more likely to persist with the previous rule after the change point (t = 2.75, p < 0.01).

#### Reinforcement-learning models

In the learning phase, the high-volatility group had a higher learning rate than the low-volatility group, as estimated by the RW-RL model 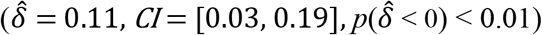. Learning rates from the 2R-RL model also showed a main effect of volatility 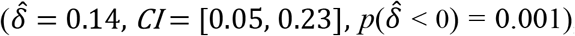, which was again driven by the high-volatility group having a higher learning rate. We additionally found a main effect of feedback 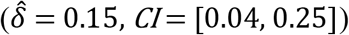, due to higher learning rates for the negative feedback trials. There was no feedback × volatility interaction 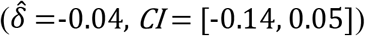.

In the transfer phase, the RW-RL model showed no significant difference of learning rates between groups 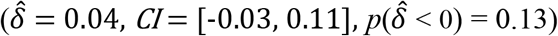, though the high-volatility group showed a numerically higher learning rate. The 2R-RL model showed no main effect of volatility 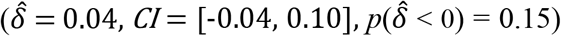, but there was a main effect of feedback 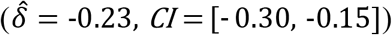, which was driven by higher learning rates for positive feedback trials. Critically, results also showed a significant feedback × volatility interaction 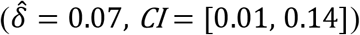. The interaction was driven by the high-volatility group showing a higher learning rate compared to the low-volatility group for negative feedback 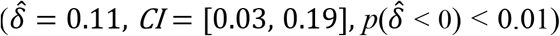, but not for positive feedback trials 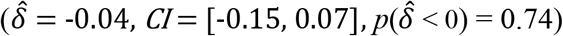. Thus, learning rates acquired during the learning phase generalized to rule switching performance in the transfer phase despite a change in specific task rules between phases, but only during negative feedback.

Even though in Experiment 2 the task rules that participants were switching between changed from the learning to the transfer phase, we observed robust evidence for transfer of rule learning rates. These results suggest that participants do not transfer a specific association between particular task rules and change point estimates in the present paradigm, but rather that they form and transfer a more abstract expectation of the volatility of the rules governing the environment, as reflected in the learning rate. This transfer was expressed primarily in response to negative feedback rather than to positive feedback, in line with the assumption that negative feedback trials in particular cause participants to switch rules.

## Experiment 3

### Method

Experiments 1 and 2 showed that task rule learning rates can generalize in time and across task rules. The cross-rule transfer effect observed in Experiment 2 clearly indicates that participants formed a rule-independent expectation of rule volatility, though transfer of learning rates here may have been aided by the fact that the new rules were still applied to the same stimuli. To further probe rule learning rate transfer to a more dissimilar environment, in Experiment 3, we provided a test of “far transfer” by testing whether prior experiences of low- or high-volatility environments would bias the tendency to shift sets in subsequent medium-volatility environments with both novel rules *and* novel stimuli.

#### Participants

One-hundred-and-one participants were recruited from MTurk and randomly assigned to one of two volatility groups. Participants were compensated at a base pay rate of $2.50 plus any additional bonuses (mean = $1.82, SD = $0.09) earned during the experiment. Twenty participants were excluded from the analysis due to overall accuracy lower than 65%, leaving a final sample size of 81. The low-volatility group had 39 participants (26 male, 12 female, 1 did not wish to reply; age range: 27-67, mean = 41.41, SD = 10.94) and the high-volatility group had 42 participants (23 male, 18 female, 1 did not wish to reply; age range: 21-56, mean = 36.29, SD = 7.88).

#### Stimuli

We used the same stimuli as in Experiments 1 and 2 for the learning phase of Experiment 3. To test whether the rule learning rates could transfer to other tasks with novel stimuli, for the Experiment 3 transfer phase we used face stimuli taken from the Chicago Face Database (Ma, Correll, & Wittenbrink, 2015). A total of 64 emotion-neutral faces (16 Asian males, 16 Asian females, 16 Caucasian males, and 16 Caucasian females) were used.

#### Procedure

In the first half of the experiment (the learning phase), participants had to sort cards according to two of the four dimensions (e.g., color and shape; randomly assigned across participants). Before the start of the second half (transfer phase) of the experiment, participants were taken to another instructions screen and informed that they would now be sorting face images according to either gender (male vs. female) or race (Asian vs. Caucasian). As in the card-matching task, on each trial three faces were displayed arranged in a pyramid, with the face on the top serving as the reference face, and the faces at the bottom as choice faces. Each of the two choice faces shared only one matching domain (gender or race) as the reference face. There was no practice for the transfer phase, and it started as soon as participants indicated they were ready.

### Results

#### Behavior

Both groups were able to perform the task reasonably well (low-volatility group: mean = 74.75%, SD = 5.66%; high-volatility group: mean = 71.96%, SD = 4.81%). The phase (learning vs. transfer) × volatility (low vs. high) ANOVA showed a significant main effect of phase (*F*(1,79) = 11.38, *p* = 0.001), a main effect of volatility (*F*(1,79) = 5.74, *p* = 0.02), and a phase × volatility interaction (*F*(1,79) = 16.28, *p* < 0.001). As in the prior two experiments, the main effect of phase was driven by higher accuracy for the transfer phase (*t* = 3.36,*p* < 0.001). In line with previous results, mean accuracy was higher for the low-volatility group in the learning phase (*t* = 4.45, *p* < 0.001), but not in the transfer phase (*t* = −1.24, *p* = 0.22).

Examination of participants’ rule choice probability before and after the rule change using a phase (learning vs. transfer) × boundary (before vs. after) × volatility (low vs. high) ANOVA showed a significant effect of boundary (F(1,79) = 685.66, p < 0.001). There were no main effects of phase (F(1,79) = 3.56, p = 0.06) or volatility (F(1,79) = 2.58, p = 0.11). There was a boundary x volatility interaction (F(1,79) = 4.10, p < 0.05). None of the other interactions were significant (all Fs(1,79) < 0.84, all ps > 0.36). We conducted separate boundary (before vs. after) × volatility (low vs. high) ANOVAs on the learning and transfer phases. In the learning phase, we find a main effect of boundary (F(1,79) = 350.51, p < 0.001), but no main effect of group (F(1,79) = 0.59, p = 0.45), and no boundary × volatility interaction (F(1,79) = 1.28, p = 0.26). Direct comparisons between volatility groups showed no difference in both before (t = −0.14, p = 0.89) or after (t = 1.18, p = 0.24) the rule change point. In the transfer phase, we find a main effect of boundary (F(1,79) = 350.15, p < 0.001), but no main effect of group (F(1,79) = 3.45, p = 0.07), or boundary × volatility interaction (F(1,79) = 3.07, p = 0.08). T-tests between volatility groups showed there were no differences between the low- and high-volatility groups (t = 0.31, p = 0.75) before the change point; however, the low-volatility group was significantly more likely to perform the previous task rule after the change point (t = 2.27, p = 0.03).

#### Extended choice probably analysis

While a full computational theory of volatility learning and the transfer thereof is beyond the scope of this paper, a closer examination of choice behavior across the task provides some evidence as to the mechanism by which transfer learning occurred. The time-course of learning shows that behavior adjusted within the first quarter of the learning phase but remained stable when environmental volatility changed at the onset of the transfer phase (Supplementary Figure 2). This suggests that the learning of the volatility level slows down over the course of the task, leading to transfer between temporally adjacent environments. This is consistent with a learning process that tracks the volatility of the environment by learning the variability of rewards in the environment but, does so with a volatility learning rate that decreases over time (Supplementary Results).

#### Reinforcement-learning model

In the learning phase, the RW-RL model found no significant difference between learning rates in the two volatility groups 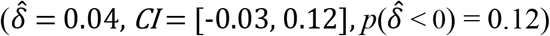, though the high-volatility group had a numerically higher learning rate compared to the low-volatility. The 2R-RL model showed the high-volatility group had higher learning rates than the low-volatility group 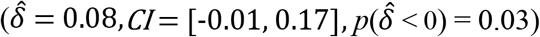. There was no main effect of feedback 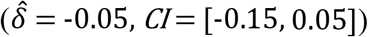. However, there was a significant feedback × volatility interaction 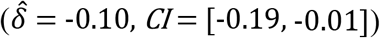. Post hoc analysis suggest that the interaction was driven by the high-volatility group showing higher learning rates than the low-volatility group for positive 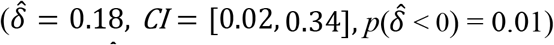 but not negative feedback 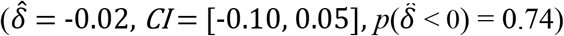.

In the transfer phase, the RW-RL model had higher learning rates in the high-volatility group compared to the low-volatility group 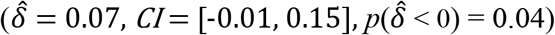. Similarly, in the 2R-RL model, the high-volatility group had higher learning rates than the low-volatility group 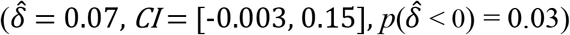. We found a main effect for feedback 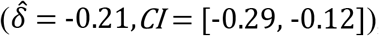, driven by higher learning rates during positive feedback trials. We found no feedback × volatility interaction 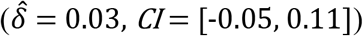. Although this interaction was not significant, based on our previous findings we further examined the volatility effect in positive and negative feedback separately, and found that the volatility effect was significant in the negative 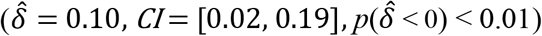, but not positive feedback trials 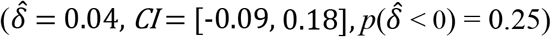. Thus, learning rates acquired during the learning phase generalized to rule switching performance in the transfer phase, in spite of a change in both the stimulus materials and task rules between phases.

In sum, we observed rule- and stimulus-independent “far transfer” of rule learning rates, in particular for negative feedback trials. In a final analysis, we sought to directly compare the degree of learning rate transfer between experiments, testing whether transfer differed quantitatively as a function of whether rules and stimuli remained the same (Experiment 1), whether rules changed (Experiment 2), or whether both rules and stimuli changed between learning and transfer phases.

#### Cross-experiment transfer effect comparison

We compared the transfer phase learning rates across the three experiments. Results from the RW-RL model showed a main effect of volatility 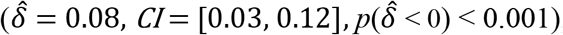, with the high-volatility groups having higher learning rates than the low-volatility groups. There were no pairwise differences in overall learning rates across the three experiments 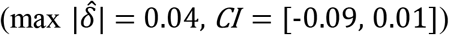. There were no interactions between volatility and experiments 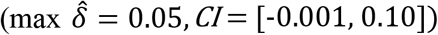.

With learning rates from the 2R-RL model, we found a main effect of volatility 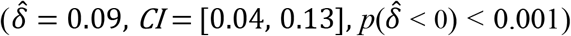, with the high-volatility groups having higher learning rates than the low-volatility groups. There was also a main effect of feedback 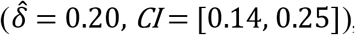, driven by higher learning rates during negative feedback trials. There were no differences in mean learning rates between pairwise comparisons across experiments 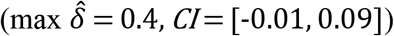. There was no volatility × feedback × experiment interaction 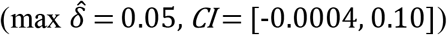.

These results echo the previous experiments in providing evidence for a transfer of task rule learning rates, even though in Experiment 3 this involved applying new rules to new stimuli (“far” transfer). Our results again also suggest that this transfer occurs primarily for negative rather than positive feedback trials. A comparison of the three experiments showed that the degree of this transfer did not differ between experiments, and was therefore unaffected by the similarity between learning and transfer tasks.

## General Discussion

The current study examined whether participants acquire and transfer expectations about cognitive flexibility demands across different contexts. We tested whether learning to change task sets more or less frequently in one context would affect learning rates in subsequent contexts. We replicated previous findings showing that participants adjusted their learning rates according to environment volatility (Behrens et al., 2007; Massi et al., 2018), with high volatility environments leading to higher learning rates, and extended that finding from stimulus-specific associations to stimulus-independent rules. Crucially, we further found that the inductive biases acquired during the learning phase affected learning rates in a subsequent transfer phase (Experiment 1) and generalized to novel task rules (Experiment 2) and novel rules and stimuli (Experiment 3). This was reflected by an overall higher learning rate in the transfer phase for participants previously exposed to a high-volatility environment compared to those previously exposed to a low-volatility environment, which was mainly driven by learning from negative feedback (unrewarded) trials. Taken together, this demonstrates that people form and transfer an abstract, stimulus- and task-independent, expectation of the volatility of the rules governing their environment, expressed in a more or less cognitively flexible rule updating strategy. To the best of our knowledge, this is the first demonstration of people’s ability to extract and re-use cognitive control learning parameters that transcend specific stimuli and tasks.

Previous behavioral studies have shown that participants can strategically adapt their readiness to switch tasks in line with changes in contextual switch-likelihood (reviewed in Braem & Egner, 2018). For instance, when manipulating the frequency of cued task switches over different blocks of trials, participants exhibit smaller switch costs (slower and more error prone responses for switching than repeating tasks) in blocks where switches are frequent compared to when they are rare (e.g., Chiu & Egner, 2017; e.g., Dreisbach & Haider, 2006; Leboe et al., 2008; Monsell & Mizon, 2006; Siqi-Liu & Egner, 2020). However, unlike in the present experiments, this change in switch-readiness seems to be limited to “biased” task sets that are associated with either more frequent switch or repeat trials, and does not generalize to intermingled “unbiased” task sets where the likelihood of switching versus repeating a task is equal (Siqi-Liu & Egner, 2020). This suggests that meta-flexibility in cued task switching is task-set or stimulus specific, rather than being due to participants developing a more global flexible cognitive strategy or processing mode that would promote switching in general (i.e., to *any* other task). This lack of transfer to other tasks also fits with cognitive training studies that demonstrated switch costs decreased over training when the same tasks are used, but substantial costs reemerged when participants are given new tasks to switch between (Sabah, Dolk, Meiran, & Dreisbach, 2019, 2021). A possible explanation underlying this dearth of transfer in prior studies could be that frequent forced (cued) switching motivates participants to keep the multiple relevant task sets in working memory, which would ease the switching between the respective tasks, but may not necessarily transfer to new tasks (Dreisbach & Fröber, 2019; Fröber, Jurczyk, & Dreisbach, 2021).

How was transfer of switch-readiness achieved in the present study? We posit that this is likely due to reliance on a different mechanism for modulating meta-flexibility. One proposal is that one’s set point on the cognitive stability-flexibility continuum can be conceptualized in terms of an “updating threshold” – the ease with which new task rule information is allowed to enter working memory (Dreisbach & Fröber, 2019; Goschke, 2003, 2013). Thus, one way to increase flexibility may be to lower this updating threshold, which in turn could increase flexibility in a more generalizable fashion (Dreisbach & Fröber, 2019). Our experiments lacked explicit instructions for when to switch; instead, people had to discover the underlying rules based on environmental feedback amidst uncertainty (Behrens et al., 2007; Niv et al., 2015; Van Eylen et al., 2011). Self-initiated switches without explicit cueing are thought to require a higher degree of disengagement from the previous task to perform the switch (Manly, Hawkins, Evans, Woldt, & Robertson, 2002; Van Eylen et al., 2011), making it less likely for participants to keep the alternative task in working memory. Studies that examined the influence of forced-choice task switching on voluntary task switches found that increasing the proportion of forced choices, in particular in combination with high switch rates, increases voluntary task switching rates (Chiu, Fröber, & Egner, 2020; Fröber & Dreisbach, 2017). Other studies showed that by rewarding switches in a prior cued task-switching phase, it is possible to increase subsequent voluntary task switching behavior (Braem, 2017), suggesting cognitively flexibility is susceptible to its recent reinforcement-learning history. Taken together, in conditions where experience is used to learn task models (Niv, 2019), meta-flexibility may be achieved by altering one’s updating threshold or the rate at which this threshold is reached (both would be observed as a difference in RL learning rate), which in turn may promote transferable effects.

As laid out in the Supplementary Results, the current transfer effects may reflect the frequency structure during the learning phase leading to participants to continue with their previous expectations, even when the environmental volatility changes (Sabah et al., 2019). Previous studies have shown that inferring hidden underlying structural forms such as the relationships between stimuli, periodicities, or cognitive maps can enable rapid generalization of behavior to new environments (Behrens et al., 2018; Halford, Bain, Maybery, & Andrews, 1998; Kemp et al., 2010; Mark et al., 2020). For instance, in Mark et al. (2020), two groups of participants learned either hexagonal or community structure graphs, and then learned a new graph with either the same or the alternate structure. The authors found that the experience with the first graph shaped prior expectations over the underlying structure on the second graph, shown by improved task performance in the group that had the correct prior structural knowledge. By analogy, it is likely that in the current study, the structure of the card sorting tasks was generalized over task rules and stimuli using similar mechanisms of applying previously learned abstracted knowledge, in this case, the updating threshold or learning rate (Baram et al., 2020). Consequently, the frequency of the switches encountered during the learning phase drove expectations and switch readiness during the transfer phase. This supports that structural knowledge acquired in the learning phase creates an inductive bias that affects how participants make environmental inferences in the transfer phase.

In conclusion, we present a novel paradigm showing that participants transfer volatility-conditioned rule learning rates to new temporal, task, and stimulus contexts. This transfer of a task- and stimulus-independent rule learning parameter represents the formation and generalization of structural task knowledge for guiding cognitive control strategies. Given that impairments in the ability to adopt a contextually appropriate level of cognitive flexibility are thought to be central to various clinical conditions (e.g., Browning et al., 2015; Manly et al., 2002; Nassar & Troiani, 2020; Van Eylen et al., 2011), this new task protocol holds promise for developing a model-based assessment of individual differences in this ability in future studies. Furthermore, learning and transfer of cognitive strategies has been a central target in applied psychology, where “brain-training” interventions have been a popular idea to help improve cognitive functioning, but have had little success at far transfer (Simons et al., 2016). Our demonstration of far transfer of cognitive flexibility settings acquired through trial-and-error learning may open the door to new, more successful approaches in this domain.

## Supporting information

Supplementary Results

## Notes

### Competing Interest Statement

The authors have declared no competing interest.

## References

Baram, A. B., Muller, T. H., Nili, H., Garvert, M. M., & Behrens, T. E. J. (2020). Entorhinal and ventromedial prefrontal cortices abstract and generalize the structure of reinforcement learning problems. Neuron. https://doi.org/10.1016/j.neuron.2020.11.024

Barraclough, D. J., Conroy, M. L., & Lee, D. (2004). Prefrontal cortex and decision making in a mixed-strategy game. Nature Neuroscience, 7(4), 404–410. https://doi.org/10.1038/nn1209

Behrens, T. E. J., Muller, T. H., Whittington, J. C. R., Mark, S., Baram, A. B., Stachenfeld, K. L., & Kurth-Nelson, Z. (2018, October 24). What Is a Cognitive Map? Organizing Knowledge for Flexible Behavior. Neuron. Cell Press. https://doi.org/10.1016/j.neuron.2018.10.002

Behrens, T. E. J., Woolrich, M. W., Walton, M. E., & Rushworth, M. F. S. (2007). Learning the value of information in an uncertain world. Nature Neuroscience, 10(9). https://doi.org/10.1038/nn1954

Berg, E. A. (1948). A simple objective technique for measuring flexibility in thinking. Journal of General Psychology, 39(1), 15–22. https://doi.org/10.1080/00221309.1948.9918159

Braem, S. (2017). Conditioning task switching behavior. Cognition, 166, 272–276. https://doi.org/10.1016/j.cognition.2017.05.037

Browning, M., Behrens, T. E., Jocham, G., O’Reilly, J. X., & Bishop, S. J. (2015). Anxious individuals have difficulty learning the causal statistics of aversive environments. Nature Neuroscience, 18(4), 590–596. https://doi.org/10.1038/nn.3961

Chiu, Y. C., & Egner, T. (2017). Cueing cognitive flexibility: Item-specific learning of switch readiness. Journal of Experimental Psychology: Human Perception and Performance, 43(12), 1950–1960. https://doi.org/10.1037/xhp0000420

Chiu, Y. C., Fröber, K., & Egner, T. (2020). Item-specific priming of voluntary task switches. Journal of Experimental Psychology. Human Perception and Performance, 46(4), 434–441. https://doi.org/10.1037/xhp0000725

Dreisbach, G., & Fröber, K. (2019). On How to Be Flexible (or Not): Modulation of the Stability-Flexibility Balance. Current Directions in Psychological Science, 28(1), 3–9. https://doi.org/10.1177/0963721418800030

Dreisbach, G., & Haider, H. (2006). Preparatory adjustment of cognitive control in the task switching paradigm. Psychonomic Bulletin and Review, 13(2), 334–338. https://doi.org/10.3758/BF03193853

Fröber, K., & Dreisbach, G. (2017). Keep flexible – Keep switching! The influence of forced task switching on voluntary task switching. Cognition, 162, 48–53. https://doi.org/10.1016/j.cognition.2017.01.024

Fröber, K., Jurczyk, V., & Dreisbach, G. (2021). Keep Flexible – Keep Switching? Boundary Conditions of the Influence of Forced Task Switching on Voluntary Task Switching. Journal of Experimental Psychology: Learning Memory and Cognition. https://doi.org/10.1037/xlm0001104

Goschke, T. (2003). Voluntary action and cognitive control from a cognitive neuroscience perspective. In Voluntary action: Brains, minds, and sociality. (pp. 49–85). Retrieved from https://psycnet.apa.org/record/2003-06267-003

Goschke, T. (2013). Volition in Action: Intentions, Control Dilemmas, and the Dynamic Regulation of Cognitive Control. In Action Science: Foundations of an Emerging Discipline (pp. 409–434). The MIT Press. https://doi.org/10.7551/mitpress/9780262018555.003.0016

Halford, G. S., Bain, J. D., Maybery, M. T., & Andrews, G. (1998). Induction of Relational Schemas: Common Processes in Reasoning and Complex Learning. Cognitive Psychology, 35(3), 201–245. https://doi.org/10.1006/cogp.1998.0679

Jiang, J., Beck, J., Heller, K., & Egner, T. (2015). An insula-frontostriatal network mediates flexible cognitive control by adaptively predicting changing control demands. Nature Communications, 6(1), 1–11. https://doi.org/10.1038/ncomms9165

Jiang, J., Heller, K., & Egner, T. (2014). Bayesian modeling of flexible cognitive control. Neuroscience and Biobehavioral Reviews. https://doi.org/10.1016/j.neubiorev.2014.06.001

Kemp, C., Goodman, N. D., & Tenenbaum, J. B. (2010). Learning to Learn Causal Models. Cognitive Science, 34(7), 1185–1243. https://doi.org/10.1111/j.1551-6709.2010.01128.x

Koch, I., Poljac, E., Müller, H., & Kiesel, A. (2018). Cognitive Structure, Flexibility, and Plasticity in Human Multitasking-An Integrative Review of Dual-Task and Task-Switching Research. https://doi.org/10.1037/bul0000144

Leboe, J. P., Wong, J., Crump, M., & Stobbe, K. (2008). Probe-specific proportion task repetition effects on switching costs. Perception and Psychophysics, 70(6), 935–945. https://doi.org/10.3758/PP.70.6.935

Lee, D., Seo, H., & Jung, M. W. (2012). Neural Basis of Reinforcement Learning and Decision Making. https://doi.org/10.1146/annurev-neuro-062111-150512

Manly, T., Hawkins, K., Evans, J., Woldt, K., & Robertson, I. H. (2002). Rehabilitation of executive function: Facilitation of effective goal management on complex tasks using periodic auditory alerts. Neuropsychologia, 40(3), 271–281. https://doi.org/10.1016/S0028-3932(01)00094-X

Mark, S., Moran, R., Parr, T., Kennerley, S. W., & Behrens, T. E. J. (2020). Transferring structural knowledge across cognitive maps in humans and models. Nature Communications, 11(1). https://doi.org/10.1038/s41467-020-18254-6

Marković, D., Goschke, T., & Kiebel, S. J. (2019). Meta-control of the exploration-exploitation dilemma emerges from probabilistic inference over a hierarchy of time scales. BioRxiv. https://doi.org/10.1101/847566

Massi, B., Donahue, C. H., & Lee, D. (2018). Volatility Facilitates Value Updating in the Prefrontal Cortex. Neuron, 99(3), 598–608.e4. https://doi.org/10.1016/j.neuron.2018.06.033

Monsell, S. (2003). Task switching. Trends in Cognitive Sciences, 7(3), 134–140. https://doi.org/10.1016/S1364-6613(03)00028-7

Monsell, S., & Mizon, G. A. (2006). Can the task-cuing paradigm measure an endogenous task-set reconfiguration process? Journal of Experimental Psychology: Human Perception and Performance, 32(3), 493–516. https://doi.org/10.1037/0096-1523.32.3.493

Nassar, M. R., & Troiani, V. (2020). The stability flexibility tradeoff and the dark side of detail. Cognitive, Affective and Behavioral Neuroscience, 1–17. https://doi.org/10.3758/s13415-020-00848-8

Niv, Y. (2019). Learning task-state representations. Nature Neuroscience, 22(10), 1544–1553. https://doi.org/10.1038/s41593-019-0470-8

Niv, Y., Daniel, R., Geana, A., Gershman, S. J., Leong, Y. C., Radulescu, A., & Wilson, R. C. (2015). Reinforcement learning in multidimensional environments relies on attention mechanisms. Journal of Neuroscience, 35(21), 8145–8157. https://doi.org/10.1523/JNEUROSCI.2978-14.2015

Rescorla, R.., & Wagner, A. (1972). A theory of Pavlovian conditioning: Variations in the effectiveness of reinforcement and nonreinforcement. In Classical conditioning II: Current research and theory (pp. 64–99).

Sabah, K., Dolk, T., Meiran, N., & Dreisbach, G. (2019). When less is more: costs and benefits of varied vs. fixed content and structure in short-term task switching training. Psychological Research, 83(7), 1531–1542. https://doi.org/10.1007/s00426-018-1006-7

Sabah, K., Dolk, T., Meiran, N., & Dreisbach, G. (2021). Enhancing task-demands disrupts learning but enhances transfer gains in short-term task-switching training. Psychological Research, 85(4), 1473–1487. https://doi.org/10.1007/S00426-020-01335-Y/FIGURES/5

Schulz, E., Franklin, N. T., & Gershman, S. J. (2020). Finding structure in multi-armed bandits. Cognitive Psychology, 119, 101261. https://doi.org/10.1016/j.cogpsych.2019.101261

Simons, D. J., Boot, W. R., Charness, N., Gathercole, S. E., Chabris, C. F., Hambrick, D. Z., & Stine-Morrow, E. A. L. (2016). Do “Brain-Training” Programs Work? Psychological Science in the Public Interest, 17(3), 103–186. https://doi.org/10.1177/1529100616661983

Siqi-Liu, A., & Egner, T. (2020). Contextual Adaptation of Cognitive Flexibility is driven by Task- and Item-Level Learning. Cognitive, Affective and Behavioral Neuroscience, 20(4), 757–782. https://doi.org/10.3758/s13415-020-00801-9

Sutton, R. S., & Barto, A. G. (1998). Reinforcement Learning: An Introduction.

Van Eylen, L., Boets, B., Steyaert, J., Evers, K., Wagemans, J., & Noens, I. (2011). Cognitive flexibility in autism spectrum disorder: Explaining the inconsistencies? Research in Autism Spectrum Disorders, 5(4), 1390–1401. https://doi.org/10.1016/j.rasd.2011.01.025

Watkins, C. J. C. H., & Dayan, P. (1992). Q-Learning (Vol. 8).

Yu, L. Q., Wilson, R. C., & Nassar, M. R. (2020). Adaptive learning is structure learning in time. Psyarxiv, 1–27. https://doi.org/10.31234/OSF.IO/R637C

