## Supplementary Results for "Transfer of learned cognitive flexibility to novel stimuli and task sets"

#### Inverse temperature results

With respect to the inverse temperature, we compared whether there were any differences between volatility groups (low vs. high), phases (learning vs. transfer), and across the three experiments. In both the RW-RL and 2R-RL models, we found a main effect of group (RW-RL model: $\hat{\delta}$ = -0.27, *CI* = [-0.55, -0.005]; 2R-RL model: $\hat{\delta}$ = -0.49, *CI* = [-0.87, -0.13]), with the high-volatility group showing lower inverse temperatures, suggesting more noise in choice policy. However, the group differences in inverse temperature only existed in the learning phase (RW-RL model: $\hat{\delta}$ = -0.46, *CI* = [-0.86, -0.04]; 2R-RL model: $\hat{\delta}$ = -0.85, *CI* = [-1.31, -0.41]) and not the transfer phase ($\hat{\delta}$ = -0.09, *CI* = [-0.48, 0.28]; 2R-RL model: $\hat{\delta}$ = -0.13, *CI* = [-0.51, 0.24]). There was a main effect of phase, with the inverse temperatures being higher in the learning phase (RW-RL model: $\hat{\delta}$ = 0.53, *CI* = [0.24, 0.81]; 2R-RL model: $\hat{\delta}$ = 0.83, *CI* = [0.65, 1.02]), indicating less noise in choice policy. Additionally, we found that Experiment 1 had higher inverse temperatures than Experiments 2 ($\hat{\delta}$ = 0.51, *CI* = [0.16, 0.86]) and Experiment 3 ($\hat{\delta}$ = 0.50, *CI* = [0.16, 0.84]) in the RW-RL model, but there were no significant differences between experiments in the 2R-RL model (max $\hat{\delta}$ = 0.37, *CI* = [-0.05, 0.80]). In the RW-RL model, we found no group × phase interactions ($\hat{\delta}$ = -0.18, *CI* = [-0.45, 0.10]) and no volatility × experiment interactions (max $|\hat{\delta}|$ = 0.15, *CI* = [-0.50, 0.20]). We found a phase × experiment interaction between Experiment 1 and Experiment 3 ($\hat{\delta}$ = -0.71, *CI* = [-1.08, -0.36]). Post hoc comparisons indicated that the interaction was a result of a significant difference between learning and transfer phases in Experiment 3 ($\hat{\delta}$ = 1.42, *CI* = [0.70, 2.18]), but not in Experiment 1 ($\hat{\delta}$ = 0.001, *CI* = [-0.67, 0.69]). We found no volatility × phase × experiment interactions (max $\hat{\delta}$ = 0.18, *CI* = [-0.59, 2.09]). In the 2R-RL model, we found a significant group × phase interaction ($\hat{\delta}$ = -0.36, *CI* = [-0.54, -0.18]), which, as mentioned above, was due to the group differences being present only in the learning but not transfer phase. There were no volatility × experiment interactions (max $|\hat{\delta}|$ = 0.10, *CI* = [-0.52, 0.33]). We found a phase × experiment interaction between all pairs of experiments (min $|\hat{\delta}|$ = 0.38, *CI* = [-0.60, -0.16]). Post hoc analyses indicated that the interaction was a result of a significant difference between learning and transfer phases in Experiment 2 ($\hat{\delta}$ = 1.58, *CI* = [1.12, 2.06]) and in Experiment 3 ($\hat{\delta}$ = 1.81, *CI* = [1.35, 2.31]), but not in Experiment 1 ($\hat{\delta}$ = 0.20, *CI* = [-0.21, 0.60]). Furthermore, the difference between phases was greater in Experiment 3 than in Experiment 2. Finally, we found a volatility × phase × experiment interaction between Experiment 1 and Experiment 2 ($\hat{\delta}$ = 0.31, *CI* = [0.09, 0.53]).

#### Parameter recovery

*RW Simulation*: We simulated 80 participants (40 in the low-volatility group, i.e., low learning rate; and 40 in the high-volatility group, i.e., high learning rate), each performing 120 trials. This is comparable to our actual sample sizes for Experiments 1~3 (n = 75, 82, 81, respectively). Just as in the transfer phase of our study, the task rule switched every 20 trials, and feedback validity was set to 80%. Choice behavior was simulated according to the softmax distribution (main text

Equation 7), with individual inverse temperature values for each agent (out of 80) sampled from a normal distribution with $\mu_\beta$ = 4.17 and $\sigma_\beta$ = 0.21, which were the fitted results from across our three experiments. In order to test our model's ability to reproduce group differences, we simulated the above 5 times, testing a range of group-level learning rates so as to cover the range of possible group differences (0.0 – 0.8). Specifically, the learning rates of individual participants in the low-volatility group were drawn from a normal distribution centered around $\mu_\alpha$ = 0.5, 0.4, 0.3, 0.2, and 0.1, with $\sigma_\alpha$ = 0.03, and the learning rates of individual participants in the high-volatility group drawn from a normal distribution of $\mu_\alpha$ = 0.5, 0.6, 0.7, 0.8, and 0.9, and $\sigma_\alpha$ = 0.03. This created a range of volatility group differences in mean learning rates, i.e., $\phi_\alpha$ = 0, 0.2, 0.4, 0.6, and 0.8. These learning rates contributed to value estimation updates via Equation 1 as described in the manuscript.

*RW Parameter Recovery:* We fit our simulated data with our RW hierarchical model. As seen in Supplementary Figure 1A, our models reliably reproduced the learning rates of both groups. The Pearson correlation coefficient between the simulated and fitted mean alphas was r > 0.99 (p < 0.001) for the low-volatility group and r = 0.99 (p < 0.01) for the high-volatility group. Pearson correlation coefficient between the simulated and fitted mean alphas differences was > .99 (p < 0.001).

*2R Simulation*: Simulations for our 2R-RL model (separate for positive and negative feedback) were essentially identical, with a few exceptions. We again sampled the learning rates for the low-volatility group from normal distributions around $\mu_\alpha$ = 0.5, 0.4, 0.3, 0.2, and 0.1, with $\sigma_\alpha$ = 0.06, and the learning rates for the high volatility group from normal distributions around $\mu_\alpha$ = 0.5, 0.6, 0.7, 0.8, and 0.9, with $\sigma_\alpha$ = 0.06. However, instead of only testing the 5 possible single learning rate group differences (0-0.8), we modeled 5 possible group differences for both positive and negative learning rates, producing 5 × 5 = 25 total simulations. The inverse temperature values were sampled from a normal distribution of mean = 4.39 and SD = 0.27 (which were again obtained from our fitted results).

*2R Parameter Recovery*: As shown in Supplementary Figure 1B, we found the Pearson correlation coefficient between the simulated and fitted mean positive alphas was r = 0.96 (p < 0.001) for the low-volatility group was r = 0.96 (p < 0.001) for the high-volatility group. The Pearson correlation coefficient between the simulated and fitted mean positive alphas differences was r = 0.98 (p < 0.001). We found the Pearson correlation between the simulated and fitted mean positive alphas was r = 0.97 (p < 0.001) for the low-volatility group was r = 0.92 (p < 0.001) for the high-volatility group. And the Pearson correlation between the simulated and fitted mean negative alpha differences was 0.99 (p < 0.001). We therefore show that our models can provide a reliable estimate of learning rate differences between groups.

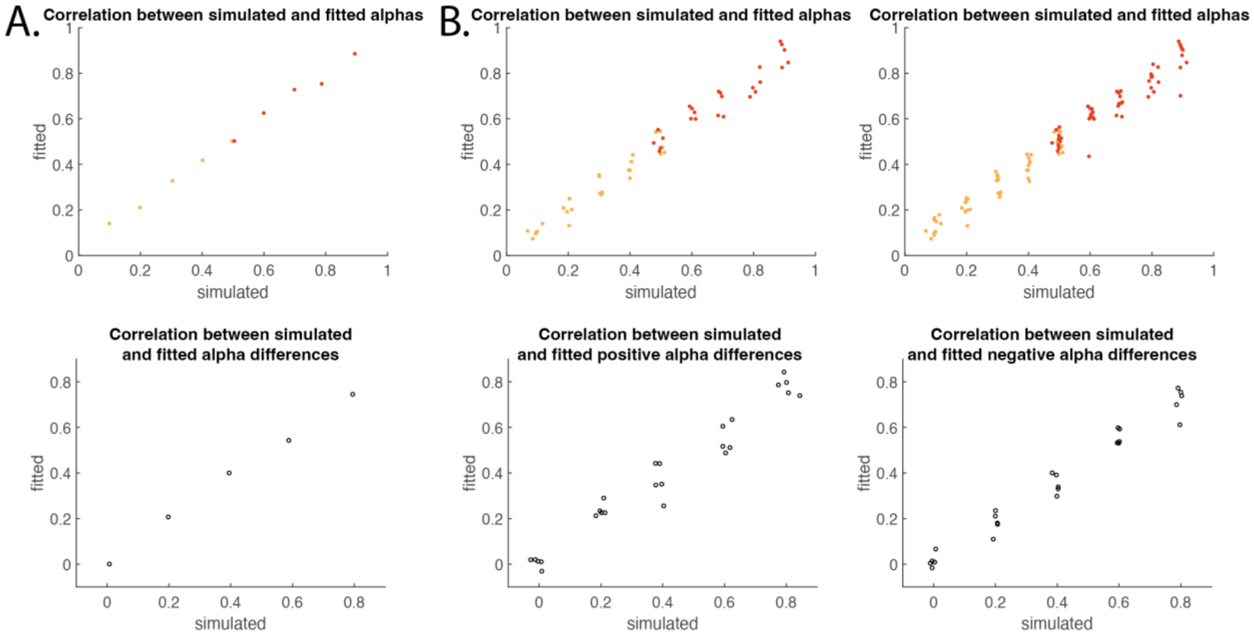


*Supplementary Figure 1. (A) Relationship between simulated and fitted alphas (top; orange: low-volatility, red: high-volatility) and alpha differences (bottom) between the high- and low-volatility groups from the RW-RL model. (B) Relationship between simulated and fitted positive (left) and negative (right) alphas (top; orange: low-volatility, red: high-volatility) and alpha differences (bottom) between the high- and low-volatility groups from the 2R-RL model.*

**Mechanisms of volatility learning**

While the present study does not allow us to identify the specific computations behind the learning and transfer of volatility expectations, a close examination of choice data can suggest possible mechanisms. For example, although we fit behavior using a simple RL model with no explicit representation of the periodic structure of the task, it is possible that participants did learn (at least approximately) that the task consisted of alternating blocks of predictable length and used this to anticipate rule changes. To look for evidence that participants anticipated switch points, we compared the probability of performing the previous rule at 1st trial after the rule change with the probability of performing the current rule 1~5 trials prior the rule change. In all experiments and conditions, the probability of picking the pre-switch rule remains stable near 0.8 through the first trial after the rule switch; only after participants have an opportunity to receive feedback does rule choice substantially change. This suggests participants don't anticipate the task switches ahead of time, using a reactive strategy rather than using the task structure to anticipate change points. Such a strategy is consistent with the simple RL mechanism we have used to describe behavior, though it is also compatible with change point learning models that only infer change points retrospectively. Simple RL models such as the one used here can be made sensitive to environmental volatility by having a parallel learning system which tracks the average magnitude of prediction errors and adjusts the reward learning rate up or down accordingly (see, e.g., Li, Schiller, Schoenbaum, Phelps, & Daw, 2011).

Next, we asked whether we could identify temporal dynamics of volatility learning that might shed light on why transfer occurred. As participants learned the volatility over time, we would expect them to change in how quickly their behavior adapts after a rule change; accordingly, their accuracy in the 1~5 trials after each rule change can be used as a proxy for their learning rate at that point in the task. In the high volatility group, post-change accuracy increases over the first three rule changes ($3^{rd}$ vs $1^{st}$ change, $t = 2.63$, $p < 0.01$), but then remains stable over the rest of the task, including after the beginning of the transfer phase. The low volatility group, on the other hand, has a similar initial post-change accuracy ($t = 0.51$, $p = 0.61$) and never significantly changes from the first rule change (max $t = 1.41$, $p > 0.15$). This indicates that an unexpected volatility level is quickly learned at the beginning of the task, yet when the volatility level changes later in the task, learning happens slowly if at all. Thus, one possibility is that transfer of volatility expectations occurs due to the failure to update volatility expectations when volatility changes, as our results show that performance remains the same as the learning phase after transitioning to the transfer phase.

Based on the above two observations, our data is consistent with a model that learns volatility by tracking prediction error magnitudes, but updates its belief about environmental volatility with a "meta" learning rate for volatility that decays over time. To demonstrate the plausibility of this mechanism for transfer learning, we simulated a learning agent with a decaying volatility learning rate performing the same task as the participants. The agent's learned volatility over the course of the task (Supplementary Figure 3) mirrors the learning trajectories of the participants (Supplementary Figure 2), with the learned volatility rising rapidly in the high volatility condition and then dropping slowly in the transfer phase, while remaining stable throughout the low volatility condition (see *Volatility learning model* below for details of the simulation).

Decaying learning rates are a standard algorithmic policy in modern machine learning, allowing agents to find global minima in value landscapes more readily (Duchi, Hazan, & Singer, 2010; Jacobs, 1988). In humans, it has been shown that learning rates decrease with the scale of prediction errors (Pearce & Hall, 1980), such that learning stabilizes over time when the environment is stable. Thus, people might reduce their learning rate of volatility learning under the expectation that volatility itself will remain stable, resulting in an inability to adjust to changes in volatility when they occur. Such a mechanism results in several interesting predictions, for example, one might expect transfer of learning rates to be reduced or not occur when the volatility is more variable during initial learning. This aligns with previous cognitive training research suggesting that practice variability being a major moderator for training transfer, specifically, increased variability during training leads participants to be more adaptable to new stimuli and task structure (Sabah, Dolk, Meiran, & Dreisbach, 2019).

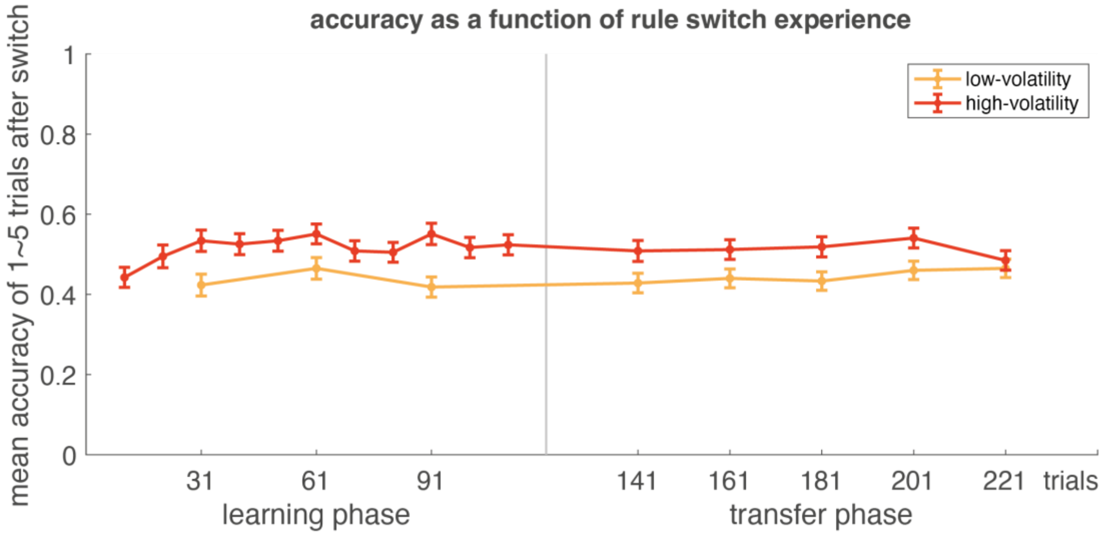


*Supplementary Figure 2. Accuracy as a function of rule change experience. Each dot represents a rule change point.*

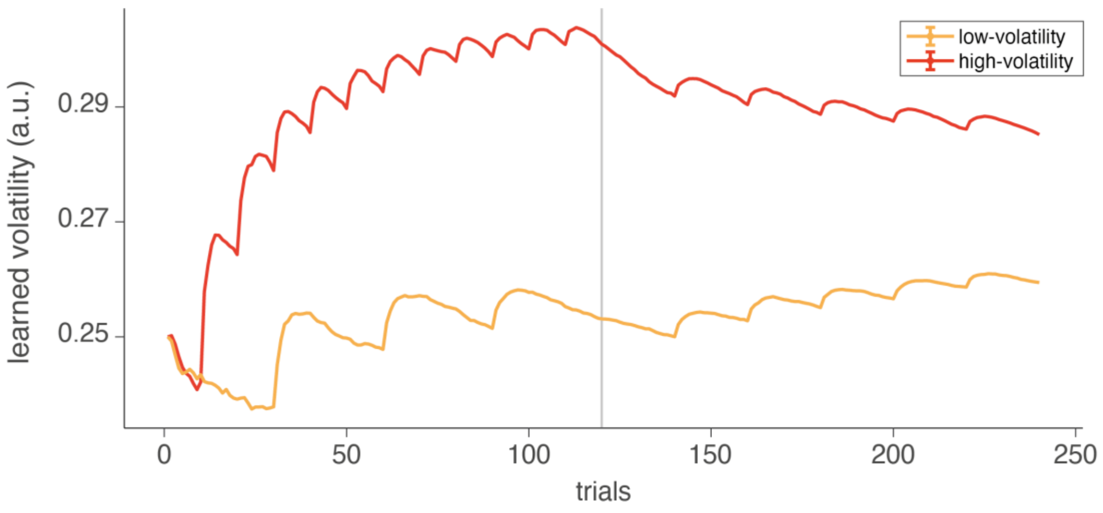


*Supplementary Figure 3. The learned volatility of simulated agents with decaying volatility learning rates. Lines represent the average of 500 simulations.*

*Volatility learning model:* To simulate the learning and transfer of environmental volatility across task phases, we simulated an agent that combined the basic RL learning method used to model behavior with a "meta" learner which tracked volatility via the variance of the prediction errors. To generate choices and track the value of each rule, the agent used equations (7) and (1) from the main text respectively, with initial starting utility and parameter values $\alpha$ and $\beta$. Rewards were generated according to same task structure experienced by participants. After each reward, the volatility $\upsilon$ was updated according to the equation:

$$\upsilon_{t+1} = \upsilon_t + \frac{\tau}{1+\omega t}(\delta^2 - \upsilon)$$

where $\delta$ is the reward prediction error from equation (1) and the decay parameters $\tau$ and $\omega$ were both fixed at 0.1. The initial expected volatility, $\upsilon_0$, was set to 0.25, to match the variance of a Bernoulli random variable with probability 0.5.

### References


Duchi, J., Hazan, E., & Singer, Y. (2010). Adaptive subgradient methods for online learning and stochastic optimization. In *COLT 2010 - The 23rd Conference on Learning Theory* (Vol. 12, pp. 257–269).

Jacobs, R. A. (1988). Increased rates of convergence through learning rate adaptation. *Neural Networks*, *1*(4), 295–307. https://doi.org/10.1016/0893-6080(88)90003-2

Li, J., Schiller, D., Schoenbaum, G., Phelps, E. A., & Daw, N. D. (2011). Differential roles of human striatum and amygdala in associative learning. *Nature Neuroscience 2011 14:10*, *14*(10), 1250–1252. https://doi.org/10.1038/nn.2904

Pearce, J. M., & Hall, G. (1980). A Model for Pavlovian Learning: Variations in the Effectiveness of Conditioned But Not of Unconditioned Stimuli. *Psychological Review*, *87*(6), 532–552.

Sabah, K., Dolk, T., Meiran, N., & Dreisbach, G. (2019). When less is more: costs and benefits of varied vs. fixed content and structure in short-term task switching training. *Psychological Research*, *83*(7), 1531–1542. https://doi.org/10.1007/S00426-018-1006-7/FIGURES/3
